# The inherent fragility of collective proliferative control

**DOI:** 10.1101/2024.01.23.576783

**Authors:** Michael G. Caldwell, Arthur D. Lander

## Abstract

Tissues achieve and maintain their sizes through active feedback, whereby cells collectively regulate proliferation and differentiation so as to facilitate homeostasis and the ability to respond to disturbances. One of the best understood feedback mechanisms—*renewal control*—achieves remarkable feats of robustness in determining and maintaining desired sizes. Yet in a variety of biologically relevant situations, we show that stochastic effects should cause rare but catastrophic failures of renewal control. We define the circumstances under which this occurs and raise the possibility such events account for important non-genetic steps in the development of cancer. We further suggest that the spontaneous stochastic reversal of these events could explain cases of cancer normalization or dormancy following treatment. Indeed, we show that the kinetics of post-treatment recurrence for many cancers are often better fit by a model of stochastic re-emergence due to loss of collective proliferative control, than by deterministic models of cancer relapse.

## INTRODUCTION

Animal species depend for survival on the ability to build and maintain tissues that are stable (relatively unchanging in time) and robust (near in size and morphology to a desired set-point). Among other things, achieving these goals requires a mechanism for proliferative homeostasis, a process typically carried out in the context of cell lineages—i.e., the production of differentiated or “terminal” cells by stem cells or stem cell-derived intermediate progenitors. Although it is widely understood that homeostasis must, at some level, amount to a process of matching the production of terminal cells to their turnover, surprisingly little is known about how this is accomplished across the multitude of tissue types.

In recent years, mathematical models have helped elucidate fundamental mechanisms underlying stem cell behavior and tissue growth^1-7^. Here we exploit such models to identify tradeoffs that tissues may encounter in attempting to achieve homeostasis. In particular, we investigate the possibility that control mechanisms that evolved to ensure tight homeostasis are also a source of fragility, so that, on very rare occasions, one may expect well-controlled tissues to spontaneously start growing without bound.

To the extent such a fragility exists—and we will argue that it is likely unavoidable—understanding how tissues minimize it can shed light on the constraints under which normal tissues operate. Perhaps even more importantly, understanding how such a fragility arises can suggest new ways to think about cancer—a situation defined by the absence of controlled growth. Cancer initiation is often portrayed of as the result of genetic changes that “drive” growth or “inhibit” cell death, but if homeostasis is controlled through feedback (and we argue here that it must be), one would not generally expect quantitative adjustments to such drives to produce a total collapse of control. In contrast, if total collapse can occur on its own at random, then an alternative way to think about oncogenes and tumor suppressor genes is as factors that make such rare transitions more likely.

Here we use simulation and mathematical analysis to show that a relative common form of homeostatic feedback control necessarily creates opportunities for rare, stochastic escape, particularly when the spatial constraints of cells and tissues are taken into consideration. At the heart of this behavior lies a stochastic transition that is collective, rather than cell-autonomous.

Although we do not currently know whether the development of some cancers depends upon the existence of this transition, we argue that its inherent reversibility could explain how some treated cancers become dormant, re-emerging stochastically at a much later time. Using clinical data on the timing of cancer recurrence in patients treated for a variety of different cancers, we show that recurrence kinetics are, indeed, often better fit by a model based on stochastic re-emergence than by traditional, deterministically motivated models.

## RESULTS

### Producing or maintaining reliable tissue size requires feedback control

Maintaining self-renewing tissues, such as blood, skin and intestinal epithelia, at fixed sizes requires stem cells to replace differentiated, or “terminal”, cells at a rate matching cell turnover. It is widely recognized that, to replace terminal cells at a constant rate, exactly half the offspring of stem cell divisions must remain stem cells—any more and a tissue will grow in size indefinitely; any fewer and it will shrink and disappear.

Less appreciated is the fact that, even if every stem cell did renew with probability exactly equal to one half, homeostasis still could not be effectively maintained. The reason is that, for most types of stem cells, division outcomes are probabilistic, meaning that sometimes cell division produces two stem cells (symmetric renewal), sometimes two differentiated cells (symmetric extinction), and sometimes one of each (asymmetric division). Even if renewal exactly balances extinction, fluctuations induced by the probabilistic nature of these outcomes will induce, in any finite-sized population, a variance in population size that grows indefinitely over time. This is illustrated in Figure 1A, which simulates scenarios that all start with the same number of stem and terminal cells, and in which each stem cell offspring differentiates independently and probabilistically, with probability of exactly one half. Although the average cell number across simulations remains roughly constant, the variance grows without bound (also see Appendix section 1).

**Figure 1.**
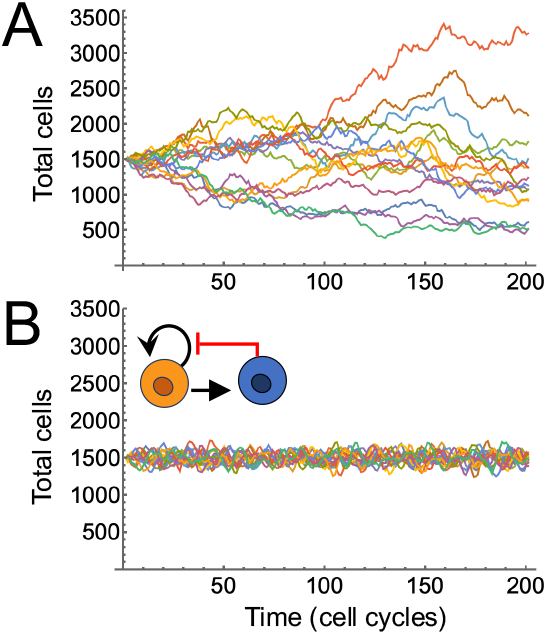
Monte Carlo simulations of stem cell systems. **A**. Starting from an initial state of 500 stem and 1000 terminal cells, stem cells divide every cell cycle, each differentiating with probability ½, and terminal cells die at a rate of 0.5 per cell cycle. Each trace is an independent simulation. **B**. Stem cells renew probabilistically, but the probability of renewal is a declining function of the number of terminal cells, with feedback strength adjusted to produce an average terminal cell number of 1000 (see Methods, eq. 1-2 and def. 1). Initial conditions and terminal cell turnover are as in panel A.

Of course, this behavior can be avoided if all stem cell divisions are asymmetric, but the fact that few stem cell systems undergo strictly asymmetric division implies that variance must get stabilized by some other means. That in turn implies that stem cells cannot behave independently of each other, a conclusion also supported by direct observation of fluctuations in systems sustained by small numbers of stem cells, e.g. ^8,9^. Specifically, stem cells must adjust their renewal behavior in response to changes in tissue size, or some proxy for tissue size, such as terminal cell number, degree of cell crowding, net rate of growth, availability of a limiting nutrient, etc. In other words, *stem cells must be feedback-controlled*.

One of the simplest and most efficient ways to achieve such control is for stem cell renewal probability to fall (i.e., for differentiation probability to rise) with the number of terminal cells, a feedback strategy that has been termed “renewal control”^5^. Renewal control has been documented in several tissues including muscle, sensory epithelia, and blood, wherein the concentrations of secreted factors (often members of the TGFβ superfamily) serve as a proxy for differentiated cell number^2,4,10-12^. Renewal control not only suppresses fluctuations (Fig. 1B), it displays a property known as “perfect robustness” or “robust perfect adaptation”^13^, whereby the steady state number of terminal cells becomes independent of all parameters that lie outside of the feedback loop itself—for example the rate of the cell cycle, the rate at which terminal cells die, or the initial number of cells that the system begins with^4,14-16^ (for the equations used to produce Fig. 1 and all subsequent figures see Methods and figure legends). Robust perfect adaptation arises because renewal control is an example of integral negative feedback, a strategy widely used in both biology and engineering to maintain systems at fixed set-points in the face of unpredictable disturbances ^17^.

Renewal control provides a simple explanation for the extraordinary capacity of self-renewing tissues to grow and regenerate to genetically specified sizes without fine-tuning of initial conditions or parameters. Renewal control can also explain the robust development of non-renewing tissues—e.g. brain, retina, cartilage, etc.—which can be modeled as instances of renewal control in which terminal cells do not turn over: As terminal cells accumulate, net stem cell renewal eventually falls below 50%, leading to stem cell exhaustion, and a static “final state” (as opposed to a steady state) where only non-dividing cells remain. As long as such a final state is much larger than the initial state, it can be shown that the final state is nearly perfectly robust to initial conditions, as well as to other parameters lying outside the feedback loop^5,14^.

Figure 2 illustrates the flexibility and robustness of renewal control in both steady– and final state contexts. Panels A and E diagram feedback circuits in which terminal cells (blue) that either do (A-D) or don’t (E-H) turn over inhibit the renewal of stem cells (orange). B and F show the solutions to (deterministic) differential equations, for a single set of parameters, in which cells are represented as continuous concentrations, and feedback follows a declining Hill function (i.e., a function that, with enough terminal cells, would drive stem cell renewal to zero); notice how steady and final states are reached, respectively, in these two cases. In panels C and G, solutions to the same equations are presented using a phase diagram, with stem and total cell numbers on the two axes, and time represented by arrows on streamlines. From these panels one can see how solutions that start from different initial conditions evolve to converge on a single fixed point (panel C) or small region on the ordinate axis (panel G). Panels D and H show the outcomes of stochastic simulations similar to those in Fig. 1—cases in which cells were treated as discrete units undergoing probabilistic divisions—and the results of 100 such simulations are plotted using phase diagrams (as in C and G). Although the stochastic and deterministic trajectories are attracted to similar locations on the phase diagram, it is clear that stochasticity allows for oscillations about the deterministic steady state, a phenomenon also apparent in Fig. 1B.

**Figure 2.**
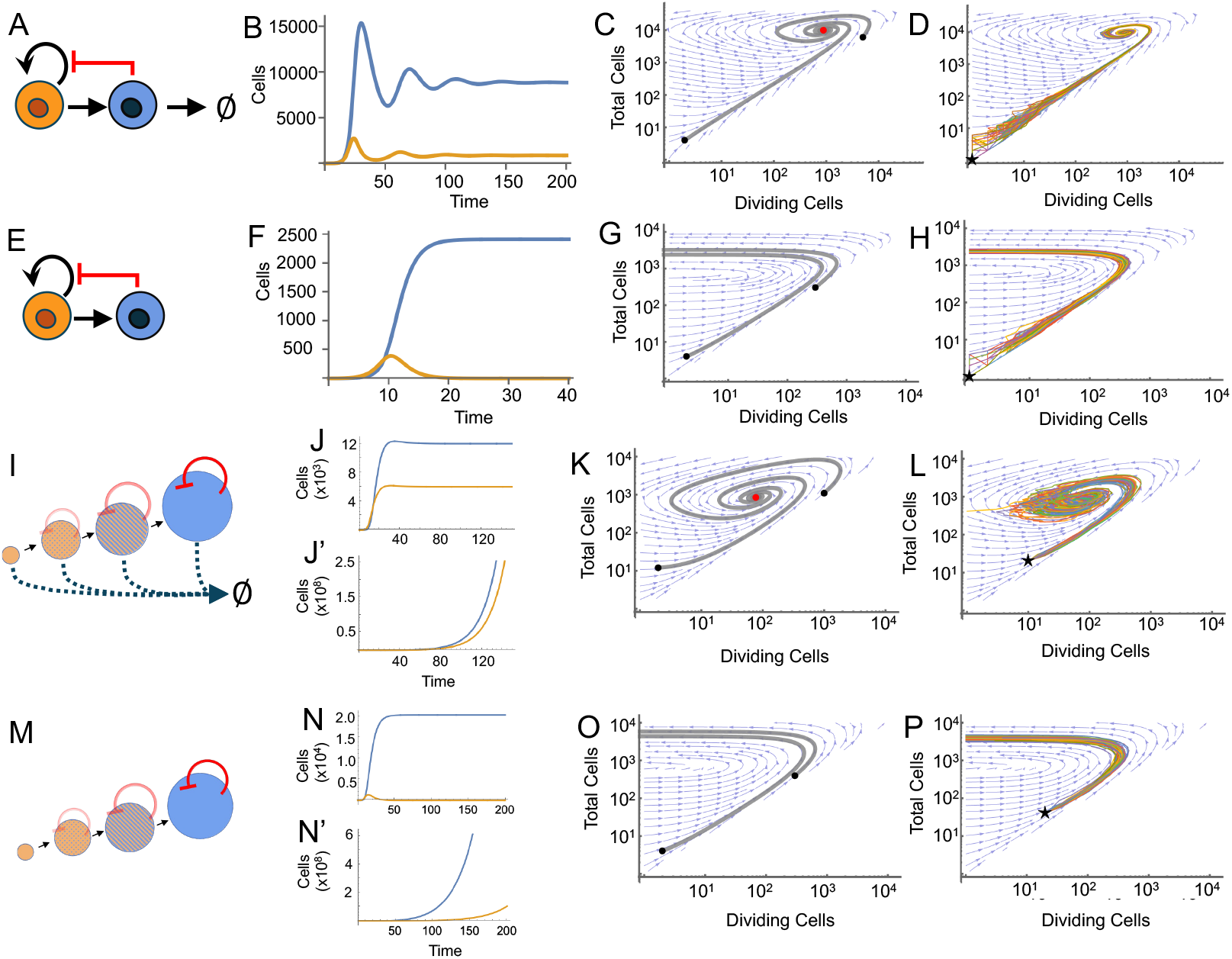
Behaviors enabled by renewal control. **A-D**. Feedback inhibition of stem cell renewal by long-lived terminal cells produces a robust steady state. Ordinary differential equations that model the feedback circuit in A were solved for a single set of parameters in B-C. In A-B, orange denotes stem cells and blue terminal cells. Panel C plots solutions in the phase plane; black dots show two sets of initial conditions, and a red dot shows the steady state. In D, 100 Monte Carlo simulations were performed in which differentiation of stem cell offspring was probabilistic; a start marks the initial conditions. **E-H**. Feedback inhibition of stem cell renewal by terminal cells that do not turn over produces a robust final state. Panels were generated and labeled as in A-D. **I-P**. Simplified models of renewal feedback in solid tissues, taking into account declining sensitivity to feedback as tissues grow beyond the characteristic decay length of a diffusing feedback signal. I-L represent the steady-state case; M-P the final-state case (no terminal cell turnover). Alternative panels J-J’ and N-N’ display stable (J, N) and unstable (J’-N’) behaviors for different parameter sets. Phase plane results are shown in K and O and stochastic simulations in L and P. For equations used, see Methods, eq. 1-2 and def. 1 (A-D) and eq. 1, 2, and 3 and def 4 (E-H). For parameter values, see Table S1.

### Stability and instability in the feedback control of growth

Any biologically useful strategy for controlling tissue size must produce a stable steady state (or, when needed, a final state). The ideal strategy should be globally stable, i.e., stable regardless of parameter choices, since that would ensure that disturbances to parameters (e.g., mutations that affect gene expression) could not easily lead to uncontrolled growth. The equations that govern the behaviors depicted in Fig. 2A-H are, in fact, globally stable. However, as with most simple models, the equations rely on making assumptions. For example, representing feedback with a declining Hill function is arbitrary. As it turns out, global stability would still be guaranteed if feedback were represented by any monotonically declining function that preserves the potential to drive the renewal probability from above ½ to below ½. A more serious issue is the assumption that the behaviors of stem cells (or the probabilities associated with those behaviors) depend only on time, and not space, i.e., stem cells all receive the same feedback regardless of location. In reality—in solid tissues at least—stem cells at different locations would likely be exposed to different levels of feedback signals. For example, if such signals are carried by diffusible molecules, concentration gradients should arise.

The stability behavior of such systems is better modeled using partial differential equations, but their analytical treatment is complicated^18^. To build intuition more easily, we considered an intermediate formulation that still uses ordinary differential equations (ODE) and accounts for spatial effects in a simplified manner. Specifically, we modeled a solid tissue as a growing disc in two-dimensions, the area of which is determined by the number of total (stem and terminal) cells. We assume stem and terminal cells are uniformly mixed (“well stirred”) within the disc (we relax this assumption later). At every location, we consider that a negative feedback signal is produced in proportion to the local concentration of terminal cells, and that the carrier of this signal diffuses away, and is taken up (or destroyed) at a constant rate. Assuming diffusion and uptake are fast relative to tissue growth, we may convert the spatial pattern of production of the feedback signal into a steady-state concentration gradient across the disc. As described elsewhere^19^, that shape may be calculated exactly, and has the expected property of being highest in the center of the disc and lowest at the edge. Accordingly, the amount of feedback “felt” by stem cells will differ by location. To obtain an expression for the aggregate behavior of the entire stem cell pool, one integrates the renewal probability across the disc, deriving an average renewal probability. The time-evolution of the total numbers of stem and terminal cells from any initial condition may then be formulated as an ODE problem.

Figure 2I-P explores the behavior of such systems for cases in which terminal cells do (Fig. 2I-L) and don’t (Fig. 2M-P) turn over. In both cases they are no longer globally stable (see Appendix section 2 for proof). When feedback is sufficiently strong, steady or final states are reached (Fig. 2J and 2N), but with weaker feedback, unbounded growth occurs (Fig. 2J’ and 2N’). This is because, as such systems expand, they experience “declining gain feedback”, where the total feedback produced by the system does not rise fast enough to keep pace with system growth. This happens because gradients created by diffusing substances produced within a spatial domain always approach a saturating shape as domain size becomes large relative to the intrinsic decay length set by the parameters of diffusivity and decay^19^.

On the other hand, such systems are still “locally stable”, meaning that, if one chooses parameters appropriately, they behave like the non-spatial models, exhibiting robust perfect adaptation of steady states and near-perfect adaptation of final states (Fig. 2K, O, respectively). Importantly, stochastic trajectories in these regimes closely mirror deterministic ones.

### Stochastic instability can arise when cell types do not thoroughly mix

Of course, solid tissues do not grow as perfect discs or spheres, nor do cells normally intermingle perfectly. To investigate the consequences of a more realistic accounting of spatial details, we turned to agent-based modeling. In such models each cell and its behavior are represented explicitly, so stochastic effects are automatically captured. Fig. 3A-B shows the rules of an agent-based model in which cells divide on a grid, pushing their neighbors aside as they do; terminal cells arise but do not turn over; and the probability that stem cell offspring renew (as opposed to differentiate) decreases according to a diffusion gradient created by the release of a feedback signal from each differentiated cell (see methods for further details). Such a model thus seeks to capture the “final state” scenario of Fig. 2M-P.

**Figure 3.**
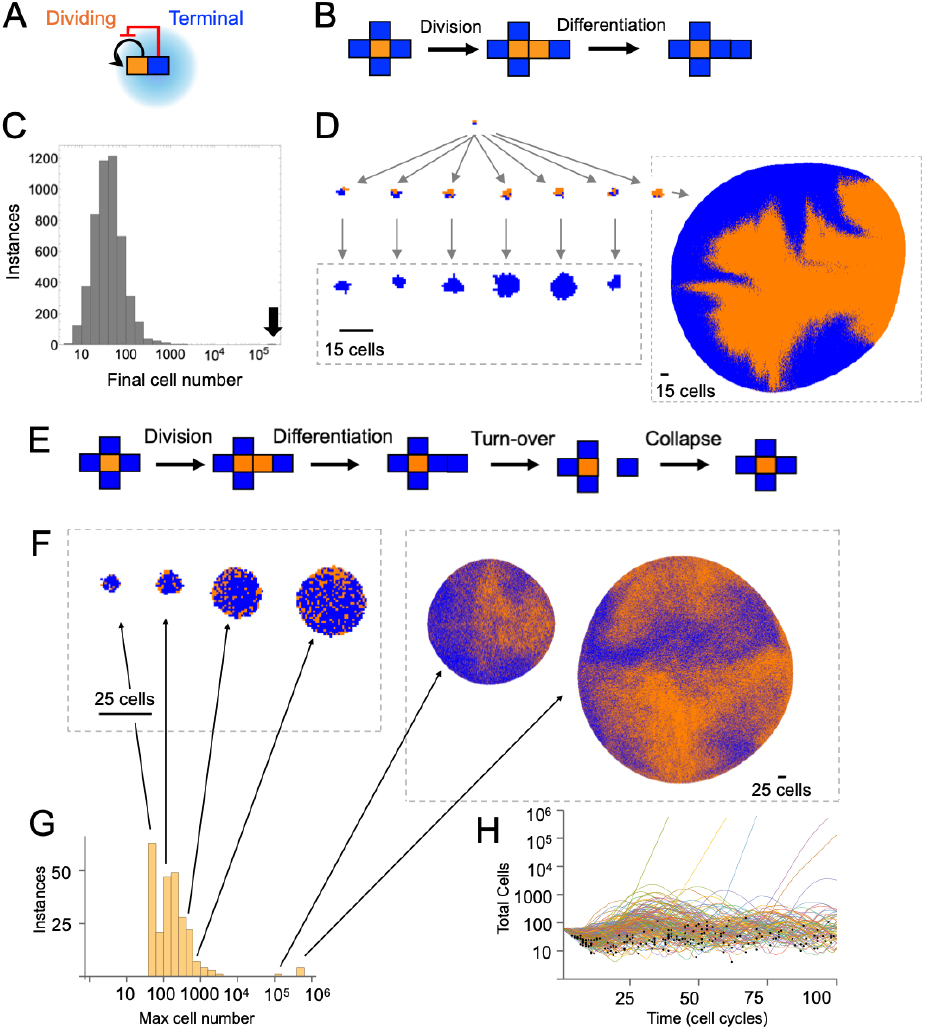
Spontaneous stochastic instability. Agent-based models were used to simulate cells growing and differentiating in a two-dimensional space, influenced by renewal feedback, either without (A-D) or with (E-H) turnover of terminal cells (see Methods). **A**. Cells were initialized on a fixed grid, and diffusion gradients from each differentiated cell calculated and summed to produce a feedback field, from which probabilities of renewal *vs*. differentiation at each position were calculated. **B**. Starting from initial conditions of two dividing cells juxtaposed with two differentiated cells, and neglecting death or removal of differentiated cells, simulations were run until either no dividing cells remained, or the number of dividing cells exceeded a pre-set maximum. **C**. Distribution of final sizes produced by 5000 such simulations. The arrow indicates rare simulations that were stopped because dividing cell numbers exceeded 70,000. **D**. Images from six representative simulations that had ceased growing, and one of three that had not. The initial condition for all simulations is shown at top, the results after 4 rounds of cell division in the middle, and those at the end of simulation at bottom. The image at right (note smaller scale bar) reached 92,998 dividing and 70,768 differentiated cells after 25 cell divisions. **E**. Starting from initial conditions close to the steady state for the chosen parameters (see Methods), simulations were run until either no dividing cells remained, the number of dividing cells exceeded 500,000, or 100 cell cycles had elapsed. **F**. Six representative simulations at their largest sizes. The four on the left continually oscillated about a steady state while the two on the right appeared to be increase approximately exponentially, and without bound. **G**. Distribution of the largest sizes (total cell number) observed among 250 simulations. The arrows show the bins into which the images in panel F fall (note logarithmic axis). **H**. Cell numbers over time for the 250 simulations summarized in panel G. Cases in which all stem cells went extinct are indicated by black dots at the time of complete stem cell disappearance. At least five cases appear to display unbounded growth (note logarithmic axis).

Simulations were initialized with a square of 4 cells, two dividing cells atop two differentiated ones, and run until either all divisions stopped, or the number of stem cells exceeded 70,000. Fig. 3C summarizes the results of 5,000 simulations, and shows that the vast majority ran out of dividing cells, stopping with a median size of 39 cells. They formed a roughly log-normal distribution of final sizes, with 99% of cases stopping at fewer than 420 cells. These simulations qualitatively mirrored the behavior in Fig. 2P, although the final size for these parameter choices was considerably smaller.

Three cases, however, continued to grow on what appeared to be an exponential trajectory. Fig. 3D contrasts the behaviors of 6 typical cases with one of the three that failed to stop growing, displaying the arrangements of cells after 4 cell cycles and, below that, at the end of the simulation.

We also modeled the scenario in which terminal cells continuously turn over so that, for sufficiently strong feedback, steady state behavior might be expected, as in Fig. 2I-L. This required modifying the agent-based simulation to allow cells to move towards each other so as to collapse empty spaces left by cell loss (Fig. 3E). Parameters were found for which simulations reached and oscillated about apparent steady states for long periods of time (e.g., 100 cell cycles), and 250 new simulations were initialized from those approximate steady state conditions. Most cases continued to oscillate about steady state, with a maximum total cell number in range of a few hundred (Fig. 3F-H). In a few cases, however, simulations switched at seemingly random times to what appeared to be unbounded exponential growth (Fig. 3H), resembling that in Fig. 3D.

Such spontaneous escape from growth control was never seen in simulations of any of the models in Figure 2. Examination of the patterns of cell arrangement in cases of escape showed that dividing and differentiated cells were no longer thoroughly mixed but had spontaneously broken up into more homogeneous patches (Fig. 3D, F). Suspecting that insufficient cell mixing might be responsible for driving rare escape from growth control, we modified the ODE models of Fig. 2 to eliminate cell mixing, by mandating that stem and terminal cells always and instantaneously sort away from each other (this allows the model to still be posed in terms of ODEs). Fig. 4A-F shows the results when stem cells were made to sort to the outside of growing discs; in Fig. 4G-L, stem cells all moved to the interior.

**Figure 4.**
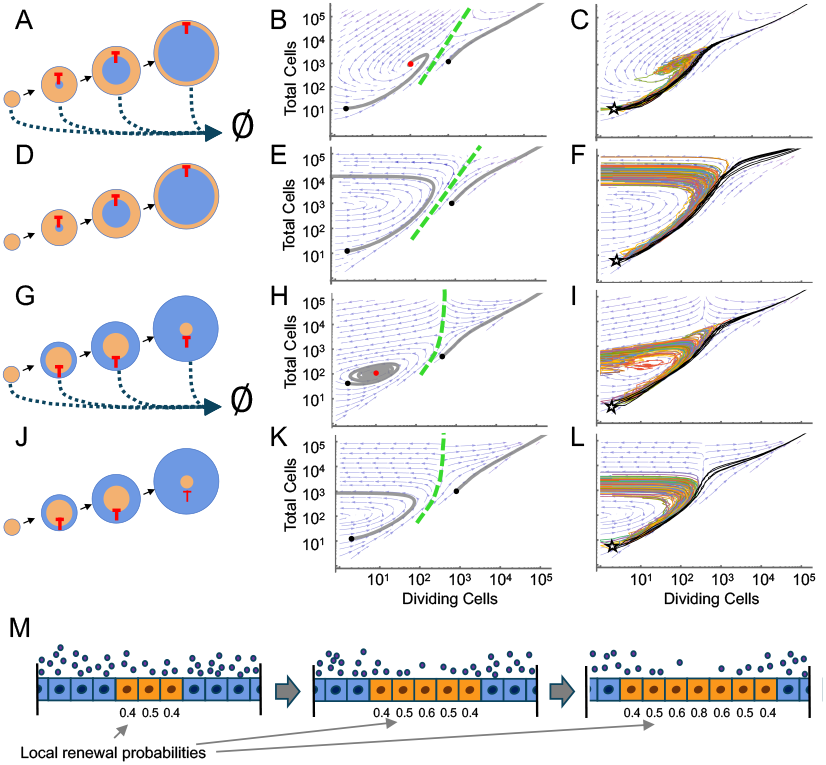
An absence of cell mixing predisposes to stochastic instability. **A**,**D**,**G**,**J**. Each row represents a system in which dividing cells (orange) grow as discs in space, and generate differentiated cells (blue), under the influence of diffusible negative feedback, modeled as in Fig. 2 panels I and M. In A and D, dividing cells instantaneously sort to the outsides of discs; in G and J they sort to the insides. In A and G, differentiated cells turn over; in D and J they do not. **B**,**E**,**H**,**K**. Phase portraits calculated for each of the four systems; note the existence of two basins of attraction in each case (separatrices are shown as dashed green lines). Black dots show initial conditions, and a red dot show steady states. **C**,**F**,**I**,**L**. 100 stochastic trajectories of each system, plotted in phase space. Initial conditions are indicated with a five-pointed star. Trajectories that spontaneously escaped from growth control are shown in black. The numbers of escaping trajectories were 7 (panel C), 8 (panel F), 6 (panel I), and 4 (panel L). **M**. Diagram explaining the source of cryptic positive feedback. If regions containing stem cells randomly grow large enough, cells in their centers will experience reduced negative feedback, favoring further growth of such regions. Equations used were eq. 1,2 and def 4 and either eq. 4 (A-F) or eq. 3 (G-L) (see Methods). See Table 1 for parameter values.

When the equations describing these scenarios were analyzed using phase diagrams (Fig. 4B,E,H,K), bimodality was noticed: For a single set of parameters, the same system could be attracted either to a stable state or undergo unbounded growth, depending on the initial conditions (depicted as black dots on the phase diagrams). In general, conditions that grew without bound were those that started with an already large number of stem cells.

Whereas initial conditions fully determined the outcomes of deterministic solutions, stochastic simulations starting from conditions expected to lead to stability occasionally ended in unbounded growth (such trajectories are shown in black in Fig. 4C,F,I and L). This is just the sort of behavior that was seen in Fig. 3. By plotting stochastic trajectories onto the models’ phase diagrams, it could be seen that escape from control arises when random fluctuations allow the system to cross, on occasion, the separatrix that divides the two possible modes of behavior.

The cartoon in Fig. 4M provides a rationale for such observations. Here stem (orange) and terminal (blue) cells are envisioned growing in one dimension, with terminal cells sorting automatically to the outside of the domain. The gradient of a diffusible feedback factor produced by terminal cells is depicted as blue dots. If, during any cell cycle, the size of the stem cell domain increases, then the average level of feedback within it will fall. The fall will be particularly large when the size of the stem cell domain exceeds the characteristic decay length of the diffusible factor. Since less feedback means more renewal, the chance that the stem cell domain will grow even larger in the next cell cycle goes up. That will lower feedback even further, further elevating renewal, until unbounded growth ultimately ensues. In effect, spatial inhomogeneity generates a cryptic positive feedback loop.

In Fig. 4, spatial inhomogeneity was enforced by the model itself (which assumes perfect cell sorting), but it is almost certainly the same phenomenon that accounts for stochastic escape in agent-based modeling (Fig. 3), where cell sorting was not mandated. This is because fluctuations in the outcomes and directionality of stem cell divisions should lead, at random, to the creation of small islands disproportionately populated with stem cells and others disproportionately populated with terminal cells. Should any one of the stem cell islands get large enough (compared with the characteristic decay length of the feedback signal), and should the average self-renewal probability within such a domain exceed ½, then effectively the same cryptic positive feedback loop as in Fig. 4 would exist.

### Positive feedback on self-renewal is sufficient to generate stochastic instability

Dynamical systems with more than one attractor state are often generated by circuits that incorporate positive feedback, sometimes combined with negative. Indeed, the behaviors of renewal control systems that utilize both positive and negative feedback on self-renewal have previously been analyzed^14^. Under appropriate conditions, such systems can be bi-stable— i.e. admitting of two stable steady states (or two possible final states).

Here we show that, under other conditions (different relative strengths of the two types of feedback), such systems can be bi-modal, producing either stability or unbounded growth, depending on the initial conditions: In Figure 5, cell dynamics were modeled using ODEs (space was not explicitly considered), and positive feedback on stem cell renewal was directly introduced, with the source either being stem cells themselves (Fig. 5A-F), or terminal cells (Fig. 5G-I). For simplicity, positive feedback was modeled as inhibition of negative feedback, although other formulations are possible^14^.

**Figure 5.**
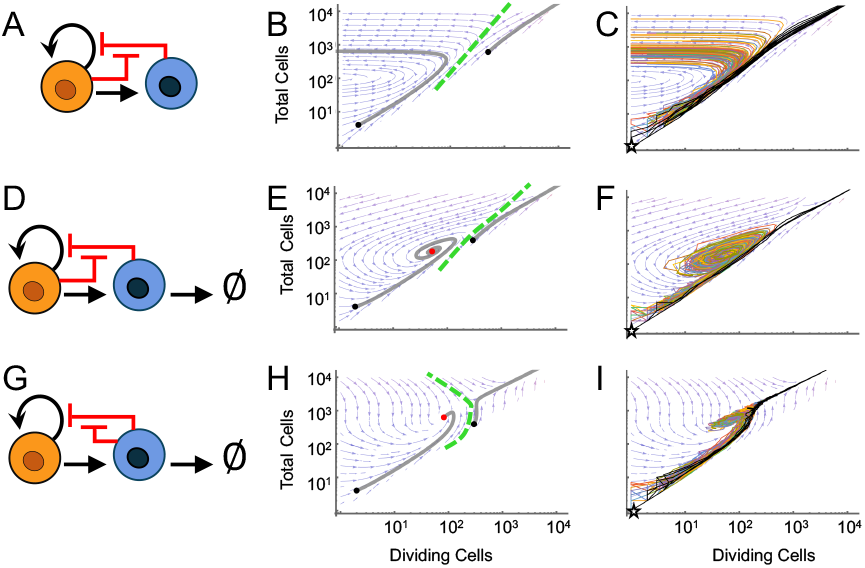
Positive feedback allows for stochastic escape from growth control. Each row presents the analysis of one of the mixed positive-negative feedback models in **A, D, and G**. Phase portraits are shown in **B, E, and H**, and stochastic trajectories (100 instances) are plotted in the same phase space in **C, F and I**. Black dots show initial conditions, and a red dot shows steady states. Initial conditions for the stochastic simulations are indicated with a five-pointed star. Trajectories that escaped growth control are shown in black. The numbers of escaping trajectories were 10 (panel C), 3 (panel F), and 6 (panel I). Equations used were eq. 1,2 and either def 2 (A-F) or def. 3 (G-I). See Table S1 for parameter values.

The results indicate that, for appropriate parameter values, mixed negative-positive feedback systems can also be bimodal, exhibiting phase diagrams much like those in Fig. 4. Moreover, stochastic simulations that started from initial conditions expected to achieve stability occasionally displayed complete loss of control, much as was also observed in Figures 3 and 4. Examination of the trajectories that escaped control suggests that, as in Figure 4, they did so because stochastic fluctuations drove them across the separatrix that divides the phase diagram into regimes of controlled and uncontrolled growth.

In summary, positive feedback on dividing cell renewal, when present, creates a scenario in which stochastic effects can produce rare escape from otherwise well-controlled growth. Such positive feedback may be an explicit process—e.g. a consequence of cell signaling—but it need not be, as effects equivalent to positive feedback tend naturally to arise spontaneously, solely due to random fluctuations in spatial homogeneity (as in Fig. 3).

### Reversibility of collective transitions

When thinking about rare, catastrophic events of unbounded cell proliferation, cancer naturally comes to mind. Although cancers have long been recognized as resulting from stochastic events, the traditional “multi-hit model” of carcinogenesis equates such events with the mutation of oncogenes or tumor suppressor genes^20,21^. Increasingly, it has been recognized that changes of a non-genetic nature, such as cells randomly flipping between epigenetic states, could also account for steps in cancer progression, and possibly also cancer initiation^22,23^. In the cancer literature, non-genetic state changes are nearly always described in cell-autonomous terms, i.e., as something that happens to a single cell, much like mutation^22^; indeed use of the term “epimutation” to describe such events has become common^24,25^.

*Collective* stochastic transitions—i.e., “tipping point” phenomena that arise out of fluctuations in collective behavior— are relatively common in organismal biology^26^, and are sometimes studied in developmental biology^27^, but have received little attention by cancer biologists (for an exception see ^28^). A unique feature of collective transitions is their propensity for reversibility in response to changes in collective circumstances. For example, if a tumor had escaped growth control through a collective transition, e.g., as modeled in Fig. 3-5, one might expect to be able to return those cells to homeostasis just by altering their numbers or spatial arrangements. In essence, if stochastic fluctuations drove crossing of a separatrix from controlled to unbounded growth, then an appropriate change of circumstances should be able to drive cells back across that same separatrix.

Simulations support this intuition. In Figure 6 the underlying model was, for convenience of analysis, the mixed positive-negative feedback model of Fig. 5G. Panel A shows 100 trajectories that all initiated from near the deterministic steady state for this system, at 2550 total cells (350 dividing and 2200 terminal). Notice how several trajectories randomly “escape” control and start to grow without bound (similar to the black trajectories in Fig. 5G). To mimic cancer treatment, we started from the point at which one of the escaped trajectories had reached 15,000 cells, and then removed a fixed number of those cells (both dividing and terminal), in the process driving the system to the other side of the separatrix shown in Fig. 5H.

**Figure 6.**
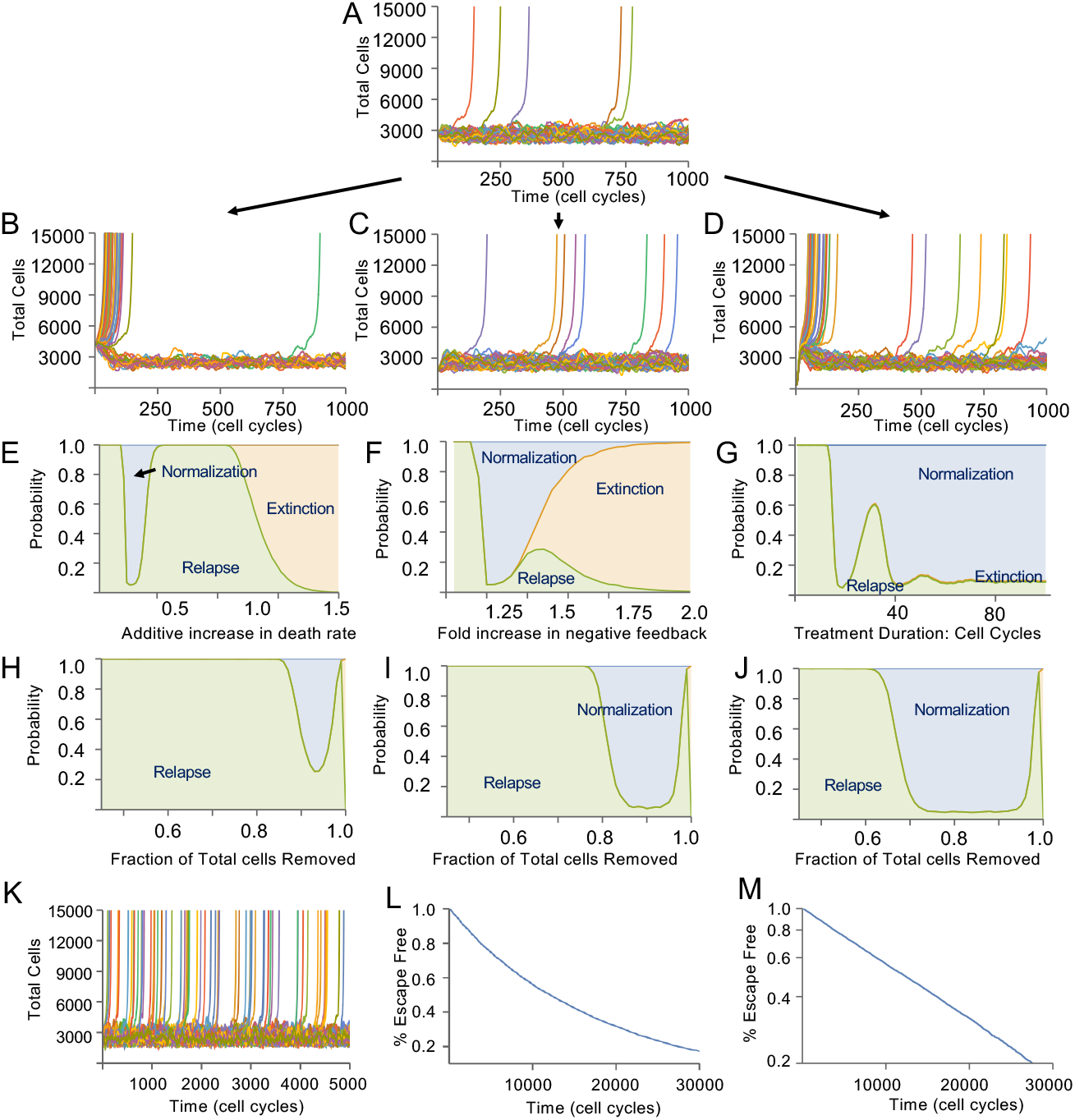
Transient treatments can reverse stochastic escape from growth control. **A**. 100 stochastic trajectories of the model in Fig. 5G initialized near steady state showing rare (∼5%) escape from growth control over the course of 1,000 cell cycles. **(B-D)** Simulations initialized above (B), somewhat below (C), and well below (D) steady state. Note the high level of early “relapse” in B and D. (**E**-**G)** Simulations were initialized from a high level (15,000 cells, of which 20% were stem cells, corresponding to the proportions observed in panel A when total cell numbers reach 15,000) and then subjected to various “treatments”, i.e., temporary or persistent changes to parameters, to determine the fraction of trajectories that normalized (returned to steady state), relapsed, or went entirely extinct (i.e., no dividing cells remaining), over the course of 1,000 additional cell cycles. In (E) treatment consisted of increasing the instantaneous rate of death of dividing and terminal cells, by the indicated amount per cell cycle, for 10 cell cycles (death was calculated *j* times per cell cycle, where *j* is the number of dividing cells). In (F) treatment consisted of increasing the strength of negative feedback by the indicated factor for 7 cell cycles. In (G) dividing cells were caused to die at an instantaneous rate of 22% per cell cycle for the indicated number of cell cycles (death was calculated *j* times per cell cycle, where *j* is the number of dividing cells). **H-J**. Simulations were initialized as in E-G, and treatment consisted of a one-time removal of the indicated fraction of total cells, (for example, a condition of 15,000 total cells and 80% removal means 3,000 remain post-treatment). The relative proportions of dividing and terminal cells removed were selected so as to produce scenarios in which the dividing cell fraction immediately after cell removal was 33.3% (H), 20% (I) or 10% (J). **K**. 250 stochastic trajectories of the model in (A) run for 5000 cell cycles, during which time 60 (24%) escaped. **L-M**. Escape-free survival (defined as never exceeding 15,000 total cells) as a function of time, for 25,000 simulations from the same initial conditions as those in K. Axes are linear in L and logarithmic in M.

Simulation was then re-initiated from initial conditions in which cell removal had driven the system to a point where total number was either above (panel B), somewhat below (panel C) or far below (panel D) the predicted (deterministic) steady state (in each case, the remaining dividing:terminal cell ratio after removal was approximately 1:6.25; see Table S1). In all cases, a large proportion of the ensuing trajectories returned to tightly controlled fluctuations, indicating resumption of homeostasis. After long times, all three conditions behaved similarly, with occasional trajectories re-escaping from control, however their short-term behaviors differed. In B, but not in C, a large proportion of trajectories returned quickly to unbounded growth. Interestingly, panel D exhibited an early burst of escaping trajectories like those in panel B. The early escapes in B undoubtedly reflect of the fact that the initial condition was sufficiently close to the separatrix that random re-crossing was relatively likely. The early escapes in D, however, reflect a different phenomenon: the inherently oscillatory nature of renewal control^5^, which causes perturbations that go too far below a steady state to elicit compensatory overshooting. Such overshoots enable a large proportion of trajectories to make close approaches to the separatrix, again allowing for a high probability of stochastic re-crossing.

The abrupt removal of a large fraction of the cells from an escaping simulation might be seen as a model of tumor treatment by surgical removal or ablation. As many cancers are treated non-surgically, e.g., with various dosing schedules of radiation, chemotherapy, or immunotherapy, we asked whether those kinds of treatments might also be expected to “normalize” the behaviors of growth-escaped cells. The results are summarized in Fig. 6E-J, in which the strength or duration of treatment is varied on the abscissa, and the ordinate shows the fraction of trajectories that either resumed homeostasis (“normalization”), fluctuated to zero (“extinction”) or exhibited early resumption of uncontrolled growth (“relapse”).

As can be seen in Fig. 6E-G, maximal normalization often occurs at relatively low treatment strengths, giving way to a high level of relapse at treatment strengths that begin to produce a large amount of extinction. Interestingly, whereas minimizing relapse with short term treatments requires fine-tuning treatment strength, it appears that low-intensity treatment over a long duration often does equally well. Although these results come from the exploration of a simplified, “toy” model, it is intriguing that they predict a phenomenon that has been observed in the clinic, which is that long-duration treatment of cancers with low dose chemotherapeutic agents (so-called “metronomic therapy”) sometimes achieves unexpectedly good results^29-32^. Results in Fig. 6H-J further show that the success of ablative therapies can depend strongly on which type of cell is primarily being preferentially targeted—dividing cells (preferentially removed in J) or their non-dividing (or slowly dividing) offspring (preferentially removed in H).

### Are there signs of stochastic instability in clinical data?

The analyses above suggest that, for cancers in which the final step in malignant transformation is a collective, stochastic transition, tumor treatment might sometimes succeed not because cells capable of initiating cancer are entirely eliminated, but because remaining malignant cells return, at least temporarily, to homeostatic growth. In fact, there are some striking examples of cancer “normalization” in animal models^47-49^, as well as rare human tumor types that spontaneously normalize without treatment^50^. There is also growing recognition that very late recurrences of tumors following treatments intended to be curative reflect “dormant” residual disease, either at the primary tumor site or elsewhere. Although mechanisms underlying dormancy are unknown^51^, it likely involves at least some degree of restoration of normal cell behavior to cells that were once cancer.

It is improbable that normalization could ever occur by chance reversion of mutations or even epimutations—both being cell-autonomous changes. Once such a change is carried by a large group of cells then, to restore homeostasis, it would be necessary for more than half of the cells to revert every cell cycle, otherwise those that didn’t would quickly replace those that did. Such reversion would be extremely unlikely, even if back-mutation (or back-epimutation) were fast at the level of the individual cell. For cancer cells that have disseminated, it has been proposed that normalization may occur because of deficiencies in the cellular environment, but exactly how that might happen, and how escape would subsequently occur, is not well understood.

With collective transitions, however, both the establishment and escape from dormancy are far easier to explain. As Fig. 6 illustrates, with systems that have escaped growth control collectively, it should be possible to re-establish a controlled steady state solely by transiently decreasing the number of cells or transiently adjusting the parameters of feedback.

Does this ever happen when cancers are treated? One way to investigate this is to examine the kinetics with which tumors recur after treatments that were intended to be curative. If, following treatment (or following metastatic dissemination), tumor cells undergo normalization through reversal of a collective transition, then the kinetics of recurrence should match those of a random process. This is illustrated by Fig. 6K-M, in which we visualize many simulated stochastic trajectories following simulated treatment (removal of a large number of cells). With sufficiently long follow-up, trajectories that had been restored to homeostasis re-escaped at times that exactly follow an exponential distribution (Fig. 6L-M). Exponential waiting times are indicative of a “single-hit” stochastic process, as they are a reflection of a probability rate (also known as a hazard rate) that is constant in time.

We wondered whether similar exponential waiting times might be observed in the recurrence of actual cancers following treatment. If so, it would strongly suggest recurrence is a “single-hit” stochastic process, consistent with a model in which tumor cells have been driven back over a separatrix across which they must once again fluctuate to escape.

We therefore gathered published clinical data from a broad range of cancers treated with curative intent (i.e., with the expectation of long-term survival of a substantial fraction of patients), focusing primarily on those in which treatment was brief and follow-up was treatment-free (i.e., we rejected cases in which recurrence could be explained by the acquisition of resistance to ongoing therapy). In such cases, the traditional explanation for recurrence is that a local or distant tumor residuum remains after treatment and continues growing until a detectable size is again reached. Even though individual residua may not all start from the same size or grow at the same rate, such an explanation is still deterministic—knowing the size and growth properties of an individual tumor residuum, one would know when that tumor would reappear. The probability rate associated with recurrence would thus not be constant in time; rather it should be peaked around the rate associated with the average tumor residuum size.

In principle, it should be possible to use clinical data on cancer recurrence to distinguish whether tumors reemerge deterministically or stochastically. In practice, however, we don’t usually know the details of how tumor residua are distributed with respect to size and growth rate, the size at which recurring tumors become recognizable, nor the fraction of patients, if any, that are cured by treatment (i.e., have no residua, and thus no probability of recurrence). Allowing these parameters to vary freely enables both stochastic and deterministic models to fit most clinical data to some degree. Still, we may investigate whether one type of model consistently does a better job of fitting across multiple data sets.

Such an analysis is shown in Figure 7 using data from 14 different clinical studies^46,49-61^. These include tumors treated by surgical resection, ablation and bone marrow transplantation, or targeted pharmacological therapy. Recurrences included both local, locoregional and metastatic. For each treated cancer in Fig. 7, the upper and middle panels compare published Kaplan-Meier curves (red) with fits to a single-hit stochastic model (blue) that allows for a detection lag (gray shaded area) and a “cured” patient fraction (orange shaded area). The middle panels show how, for most of these cancers, recurrence in at-risk patients is consistent with exponential kinetics (constant probability rate). The one clear exception is chronic myeloid leukemia (CML), in which recurrence is notably biphasic.

**Figure 7.**
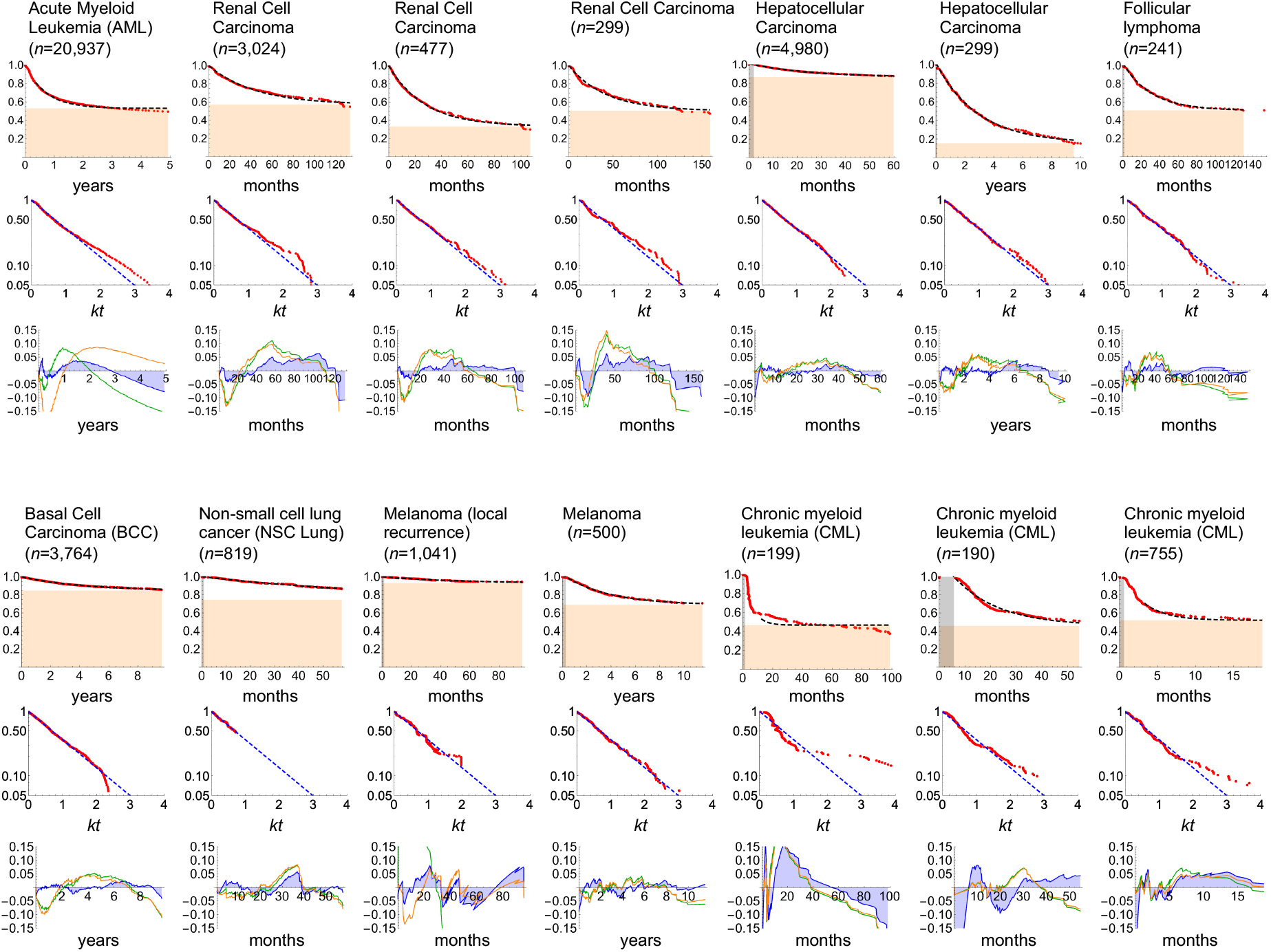
Single-hit kinetics in cancer recurrence. Reported kinetics of cancer recurrence after treatment with curative intent. For each cancer type (see Table S2 for detailed information on the data sets), the top graph reproduces a published Kaplan-Meier curve (red symbols) and blue curves represent the fit to a “single-hit” stochastic model (the fitted fraction of patients “cured” and thus not available to relapse is shown as orange fill; a thin gray bar shows the fitted detection lag, i.e. the time required before a recurring tumor becomes observable). In each middle graph, the same data were transformed to remove the influence of the cured fraction and any detection lag, so that single-hit kinetics would be expected to follow the dashed blue line of slope 1 (the abscissa shows time scaled to the fitted growth rate constant, *k*). Each lower graph plots the fit residuals for the fits in the upper graphs (the differences, over time, between the fit and the data points), not only for the single-hit stochastic model (blue curve with blue filling) but also models based on deterministic regrowth from a distribution of tumor sizes (green) or following a distribution of tumor doubling times (orange). Note that the deterministic models tend to overfit early times and underfit late ones. Measures of goodness of fit for these data are shown in Figure S3. References for these 14 studies are, in the order shown in the figure, AML^33^, renal^34^, renal^35^, renal^36^, hepatocellular^37^, hepatocellular^38^, follicular lymphoma^39^, basal cell carcinoma^40^, non-small cell lung carcinoma^41^, melanoma^42^, melanoma^43^, CML^44^, CML^45^ and CML^46^.

The bottom panels for each cancer show the fit residuals (measures of the difference between data and fit at each time point) for the stochastic model and compare them with two deterministic models that have the same number of parameters, in which tumors either start from a distribution of sizes or follow a distribution of doubling times (see Methods and Appendix section 6). The residuals of the stochastic model tended, on average, to be smaller than those of either deterministic model. More importantly, the residuals of the deterministic models tended to be negative at early times and positive at intermediate times, as expected for models with a peaked probability rate distribution. In addition, for the deterministic models, the fit parameters describing the distributions of initial residuum sizes and doubling times were often at the extremes of their allowable ranges (Supplemental Table 3).

Overall goodness of fit for the three model types was quantified in Supplemental Fig. 3A using the corrected Akaike information criterion (AICc). For all but CML, the stochastic models displayed the best AICc values. Data from an additional six studies are also presented in Supplemental Figure 3B. In some of the latter cases treatments were protracted, or we were unable to determine treatment durations, or follow-up times were very short compared to the size of the apparent at-risk population. For these reasons, confidence that such data should fit any of the proposed models was lower; nevertheless, the stochastic model performed at least as well, if not better than, other models in the majority of these cases (Fig. S3C).

## DISCUSSION

The results presented here come both from exploring theoretical models and fitting such models to clinical observations. The goal of exploring models was to build intuitions about how stochastic fluctuations and feedback control of cell proliferation might interact in unexpected ways. What was shown was that feedback circuitry that normally guarantees stability and robustness over a large portion of parameter space can, in the presence of positive feedback, also be the source of rare, stochastic instability.

The kind of positive feedback required to do this can be either explicit or cryptic. Cryptic positive feedback was shown to arise in any spatial system in which the spread of a feedback signal can in some circumstances not keep up with domain growth—at the very least this means any system in which feedback depends on diffusion, or on any other process that decays similarly over space (e.g., mechanical tension^52^). It arises particularly easily when stem cells and their differentiated offspring explicitly sort away from each other (Fig. 4) which, as it happens, is a common situation in many, if not most, epithelia, wherein stem and progenitor cells reside in a distinct compartment, usually next to the basement membrane (e.g., Fig. S2). However, as we show by agent-based modeling (Fig. 3), such a situation even arises when stem and differentiated cells simply remain near where they are born and are only moved about passively (Fig. 3).

In addition to the possibility of cryptic positive feedback due to spatial limitations, there are many tissues in which positive feedback may be explicit, through the combined actions of feedback factors that have opposing effects on self-renewal^14^. For example, tissues that use TGFβ-family molecules to feed back negatively on stem cell renewal commonly co-express MAP kinase-stimulating ligands that feed back in the opposite direction. Thus, beyond the cryptic feedback that arises due to spatial effects, there are strong reasons to expect the potential for stochastic instability to exist broadly in renewal-controlled tissues.

Stochastic instability may thus be viewed as a generic, structural shortcoming of renewal control circuitry. The idea that stabilizing, homeostatic feedback can be a source of instability may seem surprising, as control processes typically dampen, not enhance, stochastic effects (as, for example, in Fig. 1B). Such behavior nicely exemplifies what Doyle and colleagues termed the “robust-yet-fragile” nature of complex control systems—where the price of good control is typically a predisposition to rare, catastrophic failures^53^. Since the studies presented here follow from the assumption that renewal control is the mechanism that maintains homeostasis, it is possible that other classes of control mechanism might enable tissues to avoid such fragility but, as explained in Appendix section 5, short of tying the size of one tissue to another tissue, it is actually quite difficult to find other mechanisms that achieve *robust* homeostasis—i.e., maintenance of a level of terminal cells that is robust to unpredictable or uncontrollable parameters, such as the rate at which cells turnover.

Yet even if self-regulating tissues can never entirely avoid the possibility of spontaneous escape from control, one would expect them to be under evolutionary pressure to minimize the risk, which they could presumably achieve through appropriate choices of parameters (e.g., stronger negative feedback, longer decay lengths of feedback gradients). Thus, under normal circumstances, we might expect the probability of spontaneous escape from growth control during an organism’s lifetime to be kept close to zero. Oncogenic mutations, however, might change this. Indeed, thinking along these lines enables one to conceptualize oncogenes less as instructive signals that tell cells to grow, than as factors that adjust parameters so that the risk of spontaneous loss of control becomes significant. Such a view could help explain why the expression of oncogenes (or loss of expression of tumor suppressor genes) is so often associated with little or no phenotypic effect in vivo^54,55^, and why transgenic animal models with widespread oncogene expression commonly result in the production of only small numbers of tumors, e.g. ^56,57^. Whether collective, stochastic transitions to unbounded growth indeed drive cancer initiation or progression remains to be determined empirically, but several of the modeling results presented here support that view. As shown in Fig. 6, the unique tendency of collective transitions to reverse themselves following removal of cells or appropriate changes to parameters provides both a model for cancer dormancy and a justification for the ability of metronomic chemotherapy to perform comparably to high-dose, short term treatments^29-32^.

Yet by no means do these results imply all cancers should be able to revert after treatment. Even if collective transitions do play a role in carcinogenesis, subsequent cell autonomous ones (e.g., mutations and epimutations) that are effectively irreversible could block any return to normalcy. Still, the idea of reversion provides a conceptual framework for understanding cancer dormancy that does not require postulating the existence of any novel, cancer-specific mechanisms.

Clearly, the fact that cancer recurrence kinetics often fit a model in which re-emergence involves a single stochastic event (Fig. 7) does not prove the event is the one described here. Other phenomena, such as escape from immune surveillance, or induction of neovascularization, could produce similar kinetics if they occurred truly at random, but appropriate random mechanisms (e.g., mechanisms with a low, constant probability rate, such as mutation and epimutation) have not so far been found as necessary triggers of such processes. At the very least, the idea of stochastic escape from collective control provides a mechanistically plausible alternative.

The one cancer type in Figure 7 that was not well fit by the single-hit stochastic model, CML, displayed recurrence that was rapid at early times, slowing distinctly afterward. Similar bursts of early recurrence have been reported following treatment of other cancers (e.g., breast cancer), and have been suggested to reflect systemic growth-stimulatory effects triggered by surgery^58,59^. In the case of the CML, however, there is no surgical intervention; recurrence is measured following the abrupt withdrawal of a long-term suppressive pharmacotherapy, so the explanation must lie elsewhere.

Interestingly, excessive early recurrence is precisely what was seen in the simulations of Figure 6D, in which simulated tumors were reduced to a very small size by a treatment that was abruptly withdrawn. As mentioned above, the explanation for such behavior is the inherently oscillatory nature of renewal control^5^, which creates greater opportunity for stochastic escape when cell numbers start too far below their stable steady state values. In fact, in one of the CML studies included in Fig. 7, *BCR-ABL* transcripts were tracked throughout the study and frequently showed marked oscillations prior to the return of leukemia ^44^. Although such similarities do not establish definitively that CML relapse is due to collective stochastic escape, the plausibility of that explanation stands in contrast to the presumed alternative: that treated patient populations are intrinsically heterogeneous. Determining which view is correct has obvious implications for patient management.

The mathematical models of tumor recurrence that were explored here were kept deliberately simple, and the shapes of the distributions underlying deterministic models were chosen somewhat arbitrarily. Other models may well fit recurrence data equally well, especially models with greater flexibility due to a greater number of parameters—for example models in which there may be a combination of variability in both tumor size and growth rate, with or without the potential for stochastic escape. Indeed, others have generated empirical models of cancer recurrence requiring an arbitrary number of slow, sequential, stochastic events, and these also provide a good fit to clinical data^60^. Still, given that such transitions must be relatively rare to account for broad ranges of recurrence times, it is not easy to identify many possible cellular mechanisms for producing them.

Ultimately, the goal of exploring models in the present study was not so much to provide an exact match to real world data, as to provide insight into how certain kinds of behavior arise. What we can say here is that we have identified a very simple mechanism for producing random transitions to unrestrained tissue growth, a behavior that emerges spontaneously, and unexpectedly, out of the need for tissues to control their own sizes.

## Supporting information

Mathematical Appendix

## Author contributions

MC contributed to conceptualization, investigation, visualization, writing and editing. AL contributed to funding acquisition, conceptualization, supervision, investigation, visualization, writing and editing.

## Acknowledgements

This work was supported by NIH grants U54 CA217378 and P01 CA288662. MGC was supported by training grants T32 EB009418 and T32 GM136624. We thank Genti Buzi for helpful advice.

## METHODS

### Deterministic models of growth control (non-spatial)

Renewal control—in which differentiated cells dose dependently feedback upon the renewal probability of their progenitors—was adapted from^4^, which introduces the following equations:

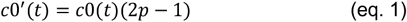

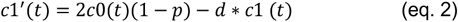

In which *c0* is the number of dividing progenitor cells, *c1* is the number of terminally differentiated cells, *d* is the turnover rate constant for differentiated cells, and *p* is the probability that a daughter cell created upon division remains dividing. If *p* is a constant, it must equal exactly ½ or the system cannot achieve a non-zero steady state. If *p* is subject to negative feedback from *c1*, i.e. is a declining function of *c1* that has a value above ½ when *c1*=0, then, for *d*>0, the system reaches a stable steady state in which dividing and differentiated (terminal) cells co-exist; if *d*=0, the system reaches a final state entirely comprised of terminal cells^14^.

For both ODE and non-spatial stochastic models, the form of the function that describes *p* differed depending on the type of feedback used into the model.

Pure negative feedback from terminal cells was represented using:

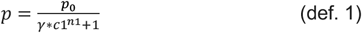

Mixed negative feedback from terminal cells and positive feedback from dividing cells was represented using:

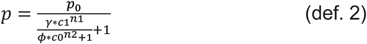

And positive and negative feedback from terminal cells was represented using:

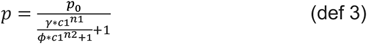

In these formulae, *p*_*0*_ is the renewal probability in the absence of feedback, *γ* and *β* are constants representing the strength of negative and positive feedback respectively, and *n1* and *n2* are the gains of negative and positive feedback respectively. Parameter values and initial conditions for all figures can be found in Table S1.

### Growing disk models

To model for the effect of space on the distribution of feedback signaling without having to solve partial differential equations, we constructed an idealized scenario in which a collection of cells was represented as a disk, the area of which was proportional to the number of cells. We then evolved discs according to equations 1-2, except the formula for *p* depended on the expected spatial distribution of a feedback signal produced by terminal cells uniformly distributed throughout the disk. Specifically, we modeled negative feedback as being carried by a factor that is produced in proportion to the number of terminal cells in the disk, that diffuses and linearly decays, reaching a steady state diffusion gradient rapidly, relative to cell growth.

In such a situation it has been shown^19^ that the steady state shape of the diffusion gradient produced is of the form

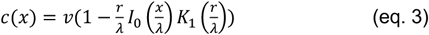

Where *c* is the concentration of the diffusing factor, *v* its rate of production, *x* the radial distance from the center of the disk, *r* the radius of the disk, *λ* a constant representing the decay length of the diffusing factor, and *I*_*_ and *K*_1_ being modified Bessel functions of the first and second kinds, respectively. To compute the renewal probability *p* for disk as whole we integrate to obtain the average of

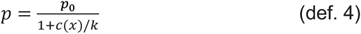

across the disk, where *p*_*0*_ is the renewal probability in the absence of feedback and *k* is a constant. In this way we obtain a single value that can be used in equations 1-2.

Under the assumption that dividing and differentiated cells were uniformly mixed within the disc, *v* was taken to be proportional to the fraction of total cells that was differentiated. In some simulations, disks were divided into an inner disk and outer annulus with dividing and differentiated cells assumed to sort immediately into one or the other compartment. In such cases it was necessary to use the formula for the diffusion gradient outside of a uniformly producing disk:

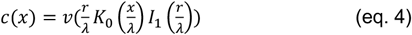

If the dividing cells occupied an annulus outside a disk of differentiated cells, Eq. 4 was used directly; if dividing cells occupied a disk inside an annulus of differentiated cells we calculated equation 3 for both the inner and outer radii and subtracted one result from the other. For parameter values see Table S1.

### Stochastic simulations of renewal control

Monte Carlo simulations were used to model the effects of fluctuations in cell number due to the probabilistic nature of cell differentiation and death. At fixed time steps all dividing cells were duplicated, and the fates of the resulting cells were probabilistically determined. Additionally, the survival of each differentiated cell was probabilistically determined, according to a constant probability of *d* of disappearing per time step. The order in which events were executed was:

1. The probability *p* of renewal was computed using the current number of dividing and differentiated cells.
2. The number of differentiated cells that die was chosen probabilistically, given *d*, and removed from the system.
3. All dividing cells were replaced by two daughter cells, and they were assigned fates so that, for any given value of *p*, the expected number of dividing cells would be *p* times the number of daughter cells. There are multiple ways in which this can be implemented, however, depending on the degree of correlation or anticorrelation in fate choice between sister cells.

The simplest approach is to have each daughter cell behave independently, differentiating with probability 1*-p*; this was the procedure used for the stochastic simulations in this study. In real-world scenarios, some stem cell populations may be inclined toward more or fewer asymmetric divisions (sisters choosing opposite fates) as opposed to symmetric ones (sisters choosing the same fate). Because asymmetric divisions balance opposing fluctuations, they tend to reduce stochastic effects. However, only when *p* is exactly ½ can all divisions be asymmetric (for any given value of *p*, the maximum proportion of divisions that can ever be asymmetric is 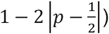, so in any system away from steady state there must always be some stochastic fluctuations.

Stochastic simulations were run until either no dividing cells remained; total cell numbers reached a predetermined limit; or a predetermined number of time steps had elapsed (see Figure legends). For parameter values see Table S1.

### Agent Based Modeling

Agent-based models were used to explore stochastic spatial effects. Conditions with the potential to reach a static final state (with no dividing cells remaining) were obtained by simulating renewal control in the absence of terminal cell death. Conditions with the potential to reach a dynamic steady state were obtained by simulating renewal control in the presence of terminal cell death.

In the first case, simulations were carried out on an 800 x 800 square grid and initiated with two adjacent dividing cells juxtaposed with two adjacent terminal cells in the center. In the second case, simulations were carried out on an 1100 x 1000 square grid, initialized from approximately steady state conditions. To identify steady state conditions, we initialized 100 simulations with a 7 x 7 grid of cells whose starting identities were chosen at random, ran simulations as described below, and identified a typical configuration characteristic of systems that remained close to a common state.

At each time step, cells could divide, and rules were used to determine both the fate of each offspring (renewal vs. differentiation; survival vs. death) and how neighboring cells would adjust their positions to the extent necessary to create an open space for a newly generated cell or close up space where a cell died. At each time step all dividing cells divided simultaneously, but the movements of other cells to accommodate new offspring were calculated sequentially.

The fate of every newborn daughter cell was assigned probabilistically, determined by the strength of local negative feedback field at its position. Field strength was determined by summing the diffusion fields produced by every terminal cell, following eq. 4, which were assumed to be of the form

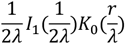

where *r* is the radial distance from the terminal cell, *λ* is a constant representing the decay length of the diffusing factor (in units of cell diameters) and *I*_1_ and *K*_*_ are modified Bessel functions of the first and second kinds, respectively. Renewal probability at each location was then calculated as

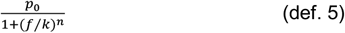

where *f* is the field strength at that location, *k* is a constant and *p*_*0*_ the renewal probability in the absence of feedback, which was usually set to 1, and *n* is a Hill coefficient set to either 1 or 2 (for steady state and final state simulations, respectively).

For final-state simulations, space for new cells was made available by identifying the nearest unoccupied grid point relative to the location of each cell that divided (if more than one location was equidistant, one was chosen at random), and then “shoving” occupied cells toward that space. “Shoving” was implemented by synchronously changing the locations of contiguous rows or columns of cells by 1 index. For example, if a cell at index (0,0) divides and the closest empty index is at (3,3), then one way to make room for a daughter cell is to free up index (0,1) by shoving the contiguous row of cells starting at (0,3) through (2,3) one index to the right (i.e., add (1,0) to each cell’s index), and then shove the contiguous column of cells starting at (0,1) through (0,2) up one index (i.e. add (0,1) to each cell’s index). Alternatively, one could free up index (1,0) by shoving the contiguous column of cells starting at (3,0) through (3,2) up one index (i.e., add (0,1) to each cell’s index) and then shove the row of cells starting at (1,0) through (2,0) one index to the right (i.e., add (1,0) to each cell’s index). These two ways of creating an unoccupied space adjacent to a dividing cell both minimize the number of cells that are ultimately shoved; for most cases there are two minimal shoving paths (row then column or column then row) and one is therefore chosen at random. When the closest unoccupied location to a dividing cell shares an index with that cell, however, only one shoving path exists. For example if the dividing cell is located at (0,0) and the unoccupied location is either (*x*,0) or (0,*x*), then only a single contiguous column or row of cells needs to be shoved. Shoving was always carried out sequentially (asynchronously) to allocate each new cell to a position as soon as it is generated. Typically, simulations were run until no dividing cells remained, or dividing cell numbers grew too numerous to simulate in reasonable times (about 70,000 dividing cells).

For simulations of cases in which terminally differentiated cells can die, it was additionally necessary to apply a constant probability of disappearance to every terminal cell and implement a rule to close up spaces created by cell death. At each time step, a random subset of differentiated cells was selected to die based on a death parameter, and subsequently, cells surrounding the unoccupied space collapsed inward to fill the void. Specifically, a cell furthest from the void was identified, and cells were shoved from that location in the manner described above, such that the void became filled. Additionally, cells divided and created additional space as already described above. Such steady-state simulations were carried out on an 1100 × 1000 square grid, initialized from approximately steady state conditions (to identify steady state conditions, we initialized 100 simulations with a 7 × 7 grid of cells whose starting identities were chosen at random, and identified a typical configuration characteristic of systems that remained close to a common state). For any given set parameters, a group of simulations was run until most fluctuated about a relative constant level of dividing and terminal cells, and this apparent steady state was used subsequently as an initial condition for the analyses shown. Simulations were then run until either all dividing cells had extinguished themselves; the number of dividing cells exceeded 500,000; or 100 cell cycles had elapsed, whichever came first.

Final state simulations were performed using *Mathematica* software; steady state simulations were hand-coded in Julia. All code will be made freely available upon request.

### Simulating tumor treatment

A non-spatial stochastic model of renewal control, which includes positive feedback but can be run as an ODE model, facilitating analysis, was used to explore how tumor treatment might influence stochastic escape from growth control. In this case the formula for the renewal probability, *p*, is that given in def. 3, which corresponds to the scenario modeled in Fig. 5G. It was determined empirically that the steady state total cell number, for the parameters chosen (Table S1), was ∼4200 and that while most simulations fluctuated above and below that number, a threshold of 15,000 total cells was sufficient to distinguish simulations that had truly escaped control, from ones that were merely fluctuating at random.

To model treatment of tumors that had collectively escaped growth control, we initialized simulations at 15,000 total cells and then, to simulate therapy, changed parameters: In the first case, we increased the rate of turnover of both dividing and differentiated cells, according to the formula

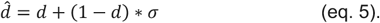

where *d* is the previous probability of death per cell per timestep (before treatment) and 0 ≤ σ ≤ 1 is a parameter representing the strength of treatment.

In a second case, treatment was implemented as an increase in negative feedback by multiplying the feedback strength *γ* by a positive constant. In a third case the feedback strength *γ* was increased by a factor of two and treatment duration was varied

In the remainder of cases, treatment consisted not in a change of parameters, but of initial conditions, corresponding to the removal of a fixed proportion of dividing cells, differentiated cells or both. The same parameters used in the previous treatment panels were used here. Treatment was begun at 15,000 total cells of which half were dividing and half differentiated. Treatment was implemented as a change in initial conditions, which were adjusted to be 15,000α(1 − σ) and 15,000(1 − α) (1 − σ), for dividing and differentiated cells, respectively, where σ (0 ≤ σ ≤ 1) quantifies overall treatment strength and α (0 ≤ α ≤ 1) quantifies any bias of treatment toward selectively removing one cell type or the other. The values of σ and *α* were chosen such that the new initial conditions would consist of dividing and differentiated cells both between 0 and 7,500. Parameters values can be found in Supplemental Table 1.

### Fitting cancer recurrence data

Cancer recurrence data were obtained from published studies (see Supplemental Table 2), in most cases by extracting data points directly from published Kaplan Meier curves (or their equivalent) and resampling in proportion to the estimated number of patients at each time point. Disease-free survival (as a fraction of total study subjects) was fit to each of three models.

<u>Model 1</u> (“stochastic emergence from dormancy”) assumes each residuum has a constant probability per time of resuming growth, expressed as *ln*2 over a rate constant *k*. Once growth has started, a delay of *ψ* must elapse before recurrence is detected.

<u>Model 2</u> (“variable tumor residuum size”) assumes the sizes of tumor residua or undetected metastases are distributed according to probability density function 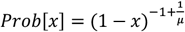, where the mean size *μ* is scaled relative to the size at which recurrence becomes detectable (i.e. *μ* =0.25 implies that the average size is one fourth of the size threshold for detection; for details see Appendix section 6). Tumors are assumed to grow with rate constant *k* (i.e., doubling time *ln*2/*k*). We constrain *μ* to be less than 0.33, as higher values would imply that the *average* residuum reaches detectable size in less than 1.6 doubling times.

<u>Model 3</u> (“variable tumor growth rates”) assumes a common tumor residuum size and tumor doubling times drawn from a log-normal distribution. The parameter *χ* characterizes the breadth of that distribution and is related to the coefficient of variation of the distribution, *c*, by 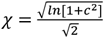 Based on clinical data we constrain to be less than 0.59 ^61^. The parameter ϕ represents the time required for a tumor residuum that starts from the average size to reach detectable size.

The parameters *ρ, k* and *ψ* (Model 1), *ρ, k* and *μ* (Model 2), or *ρ*, χ and ϕ (Model 3) were obtained by nonlinear fitting using the NonlinearModelFit function of *Mathematica* software (values are shown in supplemental Table 3).

Note that we chose to fit survival curve data directly, rather than first calculate hazard rates from data, because such calculations depend on assumptions about at-risk population sizes, and invariably create artifacts (especially at very early and late times) due to the need for data smoothing.

Note also that the values obtained for *ρ* (the “cured fraction”) were similar for all three models (see supplemental Table 3), implying that differences in quality of fit largely reflect the remaining two parameters. Note also that, for model 1, values of *ψ* were, in most cases, very small and thus had little influence on model fit. For model 2, fitted values of *μ* were, in many classes, at the maximum allowable value (0.33), implying that a better fit would have required a large number of residua to be very close to detectable size immediately after treatment, which seems implausible. Similarly, for model 3, for all cases except Chronic Myeloid Leukemia (CML) fitted values of χ were all at the maximum of 0.59, implying that a better fit would have required a greater diversity of tumor growth rates than has been observed clinically ^61^.

For more details on the derivation and limitations of the models, see Appendix section 6.

## QUANTIFICATION AND STATISTICAL ANALYSIS

When fitting cancer recurrence data, finite sample-corrected Akaike Information Criterion values, Bayesian Information Criterion values, and 95% parameter confidence intervals were obtained from the Nonlinear Model Fit object produced by *Mathematica* version 13.1.0.0. To estimate errors in AICc values, we considered all eight possible combinations of parameters set to their upper and lower confidence interval limits, and recalculated AICc using those parameter sets. The lowest absolute value of the eight results was then selected as a lower bound on |AICc|.

## SUPPLEMENTARY MATERIAL

**Figure S1. Dynamics when cells grow as a sphere in three dimensions, with spherical symmetry** (related to Figures 2 and 4). Shown are systems in which differentiated cells turn over. **A**, dividing and differentiated cells are always well mixed; **B**, differentiated cells instantaneously sort to the inside; **C**, differentiated cells instantaneously sort to the outside. Compare panels A, B, C with corresponding two-dimensional dynamics in Figures 2K, 4B and 4H, respectively. For parameter values see Table S1.

**Figure S2. Dynamics of growth control and escape from control when a planar epithelium is modeled as a flat sheet of infinite extent and variable thickness** (related to Figure 4). Growth in the apicobasal direction was modeled as in Figure 4A and G, except for the alteration to the geometry. For details see Appendix, section 3. Note that spatial sorting of dividing cells (orange) from differentiated (blue) (A) gives rise to cryptic positive feedback, leading to initial-condition dependent instability (B) and stochastic escape from control (C). For parameter values see Table S1.

**Figure S3**. Fitting cancer recurrence data sets to stochastic and deterministic models (related to Figure 7). **A**. The corrected Akaike Information Criterion (AICc)—a measure of goodness of fit—is plotted for each of the three models (single-step stochastic, variable tumor residuum size, and variable tumor residuum growth rate) for each of the cancer types analyzed in Fig. 7 (in the same order). **B**. Six additional datasets were analyzed in the manner described in Fig. 7. **C**. AICc values are shown for the three models for each of the six datasets shown in (B). In A and C, a lower error bar was obtained by calculating the 95% confidence interval around each of the three parameters in the model fits, and re-calculating AICc for every possible set of parameter values at either of the extremes of those intervals (i.e., eight pairs in total). The error bar shows the value of the AICc with the lowest magnitude, a conservative estimate of how much worse the fit quality could be if parameters were allowed to be chosen from anywhere within their confidence intervals.

**Supplemental Table 1**. Simulation parameters (related to Fig. 1-6)

**Supplemental Table 2**. Cancer data sets used (related to Fig. 7 and Fig. S3)

**Supplemental Table 3**. Parameters obtained by data fitting (related to Fig. 7 and S1).

**Supplemental Document 1**. Appendix: Mathematical derivations and model formulations (related to Fig. 1-7).

