## Supplemental figures 1-3 for "The inherent fragility of collective proliferative control"

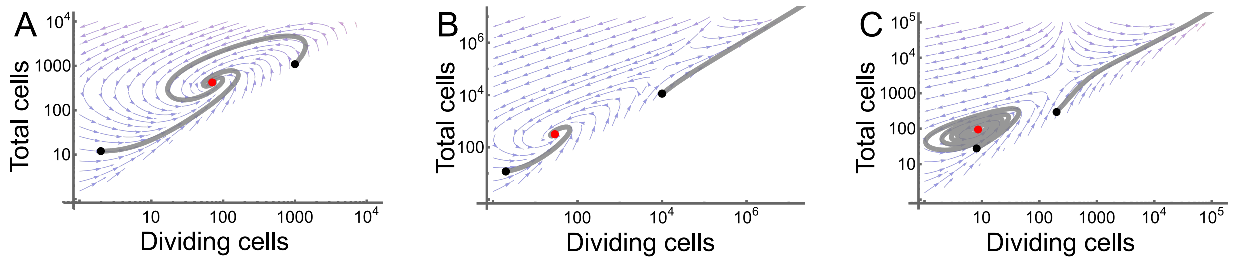


**Figure S1. Dynamics when cells grow as a sphere in three dimensions, with spherical symmetry** (related to Figures 2 and 4). Shown are systems in which differentiated cells turn over. **A**, dividing and differentiated cells are always well mixed; **B**, differentiated cells instantaneously sort to the inside; **C**, differentiated cells instantaneously sort to the outside. Compare panels A, B, C with corresponding two-dimensional dynamics in Figures 2K, 4B and 4H, respectively. For parameter values see Table S1.

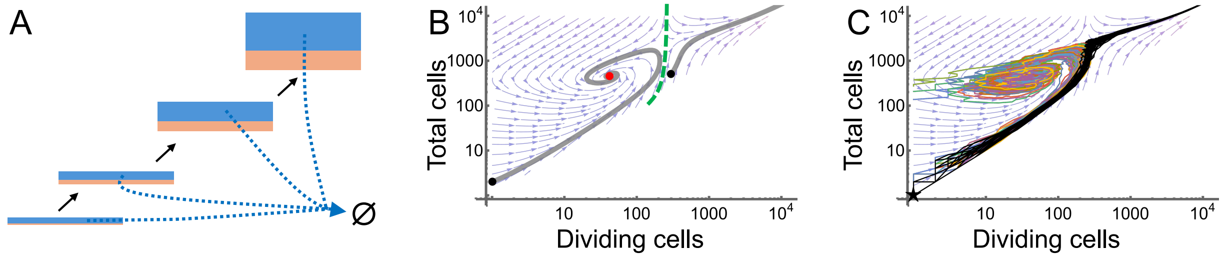


**Figure S2. Dynamics of growth control and escape from control when a planar epithelium is modeled as a flat sheet of infinite extent and variable thickness** (related to Figure 4). Growth in the apicobasal direction was modeled as in Figure 4A and G, except for the alteration to the geometry. For details see Appendix, section 3. Note that spatial sorting of dividing cells (orange) from differentiated (blue) (A) gives rise to cryptic positive feedback, leading to initial-condition dependent instability (B) and stochastic escape from control (C). For parameter values see Table S1.

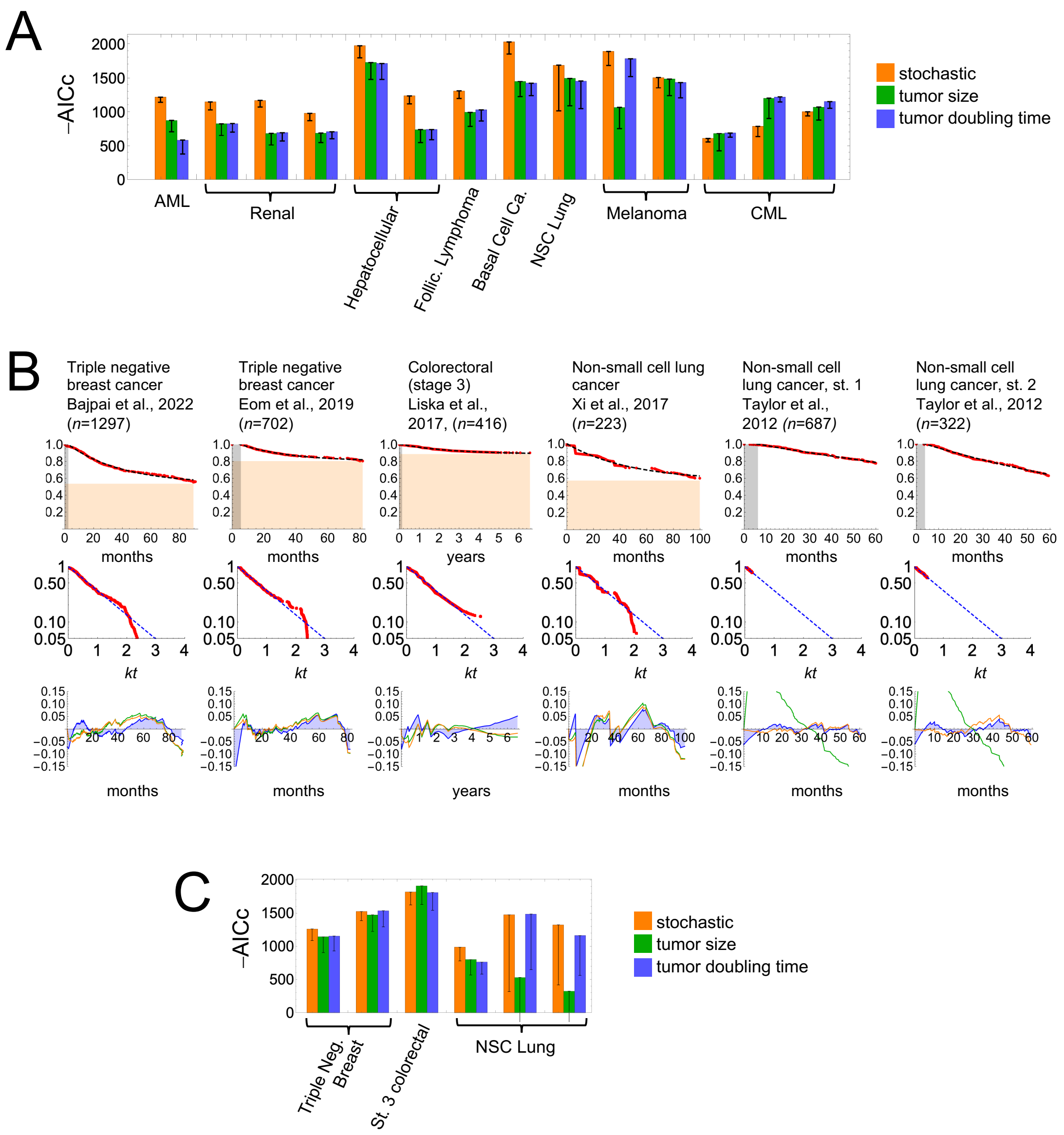


**Figure S3.** Fitting cancer recurrence data sets to stochastic and deterministic models (related to Figure 7). **A.** The corrected Akaike Information Criterion (AICc)—a measure of goodness of fit—is plotted for each of the three models (single-step stochastic, variable tumor residuum size, and variable tumor residuum growth rate) for each of the cancer types analyzed in Fig. 7 (in the same order). **B.** Six additional datasets were analyzed in the manner described in Fig. 7. **C.** AICc values are shown for the three models for each of the six datasets shown in (B). In A and C, a lower error bar was obtained by calculating the 95% confidence interval around each of the three parameters in the model fits, and re-calculating AICc for every possible set of parameter values at either of the extremes of those intervals (i.e., eight pairs in total). The error bar shows the value of the AICc with the lowest magnitude, a conservative estimate of how much worse the fit quality could be if parameters were allowed to be chosen from anywhere within their confidence intervals.
