## Supplemental table 1 for "The inherent fragility of collective proliferative control"

**Table S1.** Parameter values used in simulations (n/a = not applicable).

| Figure panel | *p0* | γ or *k* | $\phi$ | *n* (or *n1*) | *n2* | *d* | *c0* initial conditions | *c1* initial conditions | $\lambda$ | $\lambda$ /cell diameter |
|---|---|---|---|---|---|---|---|---|---|---|
| 1A | 0.5 | n/a | n/a | n/a | n/a | 0.5 | 500 | 1000 | n/a | n/a |
| 1B | 1 | $\gamma = 10^{-3}$ | | 1 | | | 500 | 1000 | | |
| 2B | | | | | | | 1 | 0 | | |
| 2C | 0.7 | γ = 0.000045 | | | | 0.1 | 2 & 5000 | 2 & 1000 | | |
| 2D | | | | | | | 1 | 0 | n/a | n/a |
| 2F | | | | | | | 1 | 0 | | |
| 2G | 0.85 | $\gamma = 10^{-3}$ | | | | 0 | 2 & 300 | 2 & 2 | | |
| 2H | | | | | | | 1 | 0 | | |
| 2J | | γ =0.7 | n/a | 1 | n/a | 0.5 | 1 | 0 | | |
| 2J' | | γ =0.9 | | | | | 1 | 0 | | |
| 2K | | γ =0.5 | | | | 0.1 | 2 & 1000 | 10 & 100 | | |
| 2L | 0.9 | | | | | | 10 | 10 | 100 | 10 |
| 2N | | γ =0.75 | | | | | 1 | 0 | | |
| 2N' | | γ =1.3 | | | | 0 | 1 | 0 | | |
| 2O | | γ =0.5 | | | | | 2 & 300 | 2 & 100 | | |
| 2P | | | | | | | | | | |
| 3C | 1.0 | *k* = 0.14 | n/a | 2 | n/a | 0 | 2* | 2 * | n/a | 2 |
| 3G | | *k* =0.263 | | 1 | | 0.25 | 9* | 49* | | 1.5 |
| 4B | | *k* = 0.46 | | | | 0.1 | 2 & 1000 | 10 & 200 | | 10 |
| 4C | | | | | | | 2 | 10 | | |
| 4E | | *k* = 0.51 | | | | 0 | 2 & 800 | 10 & 200 | | |
| 4F | 0.9 | | n/a | 1 | n/a | | 2 | 2 | 22.5 | |
| 4H | | *k* = 0.459 | | | | 0.1 | 2 & 450 | 40 & 50 | | 4.5 |
| 4I | | | | | | | 2 | 2 | | |
| 4K | | *k* = 0.52 | | | | 0 | 2 & 800 | 10 & 200 | | |
| 4L | | | | | | | 2 | 2 | | |
| 5B | | | 0.0022 | | | 0 | 2 & 500 | 2 & 100 | | |
| 5C | | γ = 0.006 | | | 1.5 | | 1 | 0 | | |
| 5E | 0.8 | | 0.00085 | 1 | | 0.4 | 2 & 300 | 2 & 100 | n/a | n/a |
| 5F | | | | | | | 1 | 0 | | |
| 5H | | γ = 0.004 | 0.001 | | 1.25 | 0.15 | 2 & 300 | 2 & 100 | | |
| 5I | | | | | | | 1 | 0 | | |
| 6A | | | | | | | 350 | 2200 | | |
| 6B | | | | | | | 575 | 3600 | | |
| 6C | | | | | | | 250 | 1560 | | |
| 6D | | | | | | | 50 | 315 | | |
| 6E | | | | | | | | | | |
| 6F | 0.9 | γ = 0.0009 | $1.41493 \times 10^{-5}$ | 1 | 1.5 | 0.15 | 3000 | 12000 | n/a | n/a |
| 6G | | | | | | | | | | |
| 6H | | | | | | | | | | |
| 6I | | | | | | | As shown | As shown | | |
| 6J | | | | | | | | | | |
| 6K | | | | | | | 350 | 2200 | | |
| S1A | 0.85 | *k* = 0.4 | | | | 0.2 | $2 \& 10^3$ | 10 & 100 | 22.5 | 2.25 |
| S1B | 0.9 | *k* = 0.8 | n/a | n/a | n/a | 0.1 | $2 \& 10^4$ | 10 & 1000 | 10.5 | 1.05 |
| S1C | 0.8 | *k* = 0.7 | | | | 0.1 | 8 & 200 | 20 & 100 | 10.5 | 1.05 |
| S2B | 0.85 | γ =1.8 | n/a | 1 | n/a | 0.1 | 1 & 300 | 1 & 200 | ** | ** |
| S2C | | | | | | | | | | |

*For Figure 3C, dividing cells were initially placed at points {400, 400} and {400, 401} on an 800 x 800 grid, and differentiated cells at points {401, 400} and {401,401}. For Figure 3G, dividing cells were initially placed at points {543,548}, {544,546}, {545,547}, {547,549}, {548,543}, {548,550}, {549,549}, {551,545}, and {552,548} on an 1100 x 1100 grid, and differentiated cells were placed at points {543,545}, {544,547}, {544,548}, {544,549}, {545,545},

{545,546}, {545,548}, {545,549}, {545,550}, {546,544}, {546,545}, {546,546}, {546,547}, {546,548}, {546,549}, {546,550}, {547,543}, {547,544}, {547,545}, {547,546}, {547,547}, {547,551}, {548,544}, {548,545}, {548,546}, {548,547}, {548,548}, {548,549}, {548,551}, {549,543}, {549,544}, {549,545}, {549,546}, {549,547}, {549,548}, {549,550}, {549,551}, {550,544}, {550,545}, {550,546}, {550,547}, {550,548}, {550,549}, {550,550}, {551,546}, {551,547}, {551,548}, {551,549}, and {552,547}. For figure 3G, per simulation time step, only 80% of dividing cells were selected to divide (i.e., the time steps represent somewhat less than 1 cell cycle).

**Here decay length was captured using a lumped constant, $\kappa = \frac{w\lambda}{\pi\sigma^2}$ , where $\lambda$ is the decay length, $\sigma$ the cell radius, and *w* is the planar width of the epithelium, i.e. the planar area of epithelium being modeled is $\pi w^2$ . In Figure S2, $\kappa = 500$.
