## Supplemental table 2 for "The inherent fragility of collective proliferative control"

**TABLE 2: Sources of cancer recurrence data for Figures 7 and Supplemental Figure 1.** Part A presents the datasets used in Figure 7D-E; part B those used in supplemental Fig. 1. Study populations sizes refer only to those patients corresponding to the Kaplan-Meier curves used in the figures.

| A. Cancer Type | Prior treatment | Criterion | Study Population Size | Citation |
|---|---|---|---|---|
| Acute myeloid leukemia | Hematopoietic stem cell transplantation | Leukemia free survival | 20,937 | (Ruggeri et al., 2016) |
| Renal cell carcinoma | Surgery ~90% of which was Radical Nephrectomy | Disease free survival, post-surgery | 3,024 | (Marconi et al., 2021) |
| Renal cell carcinoma | Nephrectomy | Overall survival, post nephrectomy | 477 | (Zisman et al., 2001) |
| Renal cell carcinoma | Nephrectomy | Disease-free survival, post-surgery | 299 | (Brookman-Amissah et al., 2009) |
| Hepatocellular Carcinoma | Liver transplant | Recurrence free survival, post-transplant | 4,980 | (Tran et al., 2023) |
| Hepatocellular Carcinoma | Hepatic resection | Disease free survival, post-surgery | 241 | (Sakon et al., 2000) |
| Follicular Lymphoma | Autologous stem cell transplantation; patients presenting with high grade transformation were excluded | Relapse-free survival | 241 | (Kornacker et al., 2009) |
| Primary Basal Cell Carcinoma | Curettage Electrodesiccation, Surgical Excision, or X-Ray Therapy | Disease-free survival, post-treatment | 3,764 | (Silverman et al., 1991) |
| Lung, non-small cell | Surgical resection plus five years follow-up without treatment or recurrence | Locoregional and distant recurrence-free survival, starting five years post-surgery | 819 | (Maeda et al., 2010) |
| Melanoma | Surgical excision | Local recurrence-free survival, post-excision | 1,041 | (Zalaudek et al., 2003) |

| | | | | |
|---|---|---|---|---|
| Melanoma | Surgical excision, observation only | Recurrence-free survival, post-excision | 500 | (Morton et al., 2014) |
| Chronic myeloid leukemia (CML) (STIM2 trial) | Imatinib | Molecular recurrence-free survival after discontinuation of therapy | 199 | (Dulucq et al., 2022) |
| Chronic myeloid leukemia (CML) (ENESTfreedom trial) | Nilotinib | Treatment-free remission after discontinuation of therapy | 190 | (Ureshino, 2021) |
| Chronic myeloid leukemia (CML) (EURO-SKI trial) | Tyrosine kinase inhibitor therapy | Molecular recurrence-free remission after discontinuation of therapy | 755 | (Saussele et al., 2018) |

| B. Cancer Type | Prior treatment | Criterion | Study Population Size | Citation |
|---|---|---|---|---|
| Breast Cancer--triple negative 1 | ~95% surgical removal<br>~97% Chemotherapy: adjuvant chemotherapy or neoadjuvant chemotherapy<br>~80% Adjuvent radiotherapy | Disease-free survival (beginning at date of diagnosis) | 1,297 | (Bajpai et al., 2022) |
| Breast Cancer--triple negative 3 | 100% Surgery<br>~78% adjuvant chemotherapy<br>~11% neoadjuvant chemotherapy | Incidence of recurrence (disease-free survival start time not specified—presumably post surgery and during chemotherapy) | 702 | (Eom et al., 2019) |
| Colorectal cancer, stage 3 | 100% Surgical removal<br>~74% Adjuvant Chemotherapy | Loco-regional recurrence free survival | 416 | (Liska et al., 2017) |
| Lung, non-small cell | 100% Surgery<br>~56% adjuvant chemotherapy. Stroma-poor tumors only, stages I, II and III. | Disease-free survival post-surgery | 223 | (Xi et al., 2017) |
| Lung, non-small cell, stage 1 | 100% Surgery, with or without induction chemotherapy or radiation | Recurrence-free survival post-surgery | 687 | (Taylor et al., 2012) |

| | | | | |
|---|---|---|---|---|
| | ~6-7% adjuvant chemotherapy | | | |
| Lung, non-small cell, stage 2 | 100% Surgery, with or without induction chemotherapy or radiation ~6-7% adjuvant chemotherapy | Recurrence-free survival post-surgery | 322 | (Taylor et al., 2012) |
