## Supplemental table 3 for "The inherent fragility of collective proliferative control"

**Supplemental Table 3**: Parameters value and measure of quality of fit for three different models of tumor recurrence. Recurrence data were extracted from the cited publications and are shown in the indicated figures. Data were normalized and fit to each of three models, "stochastic emergence from dormancy" (model 1), "variable tumor residuum size" (model 2) and "variable tumor growth rate" (model 3). Quality of fit was quantified using both AICc (finite sample corrected Akaike information criterion) and BIC (Bayesian information criterion). For meanings of model parameters, see Methods and Appendix.

| | | | Goodness of Fit (AICc) | | | Goodness of Fit (BICc) | | | Model 1 parameters | | | Model 2 parameters | | | Model 3 parameters | | | Range of $\rho$ across the three models | |
|---|---|---|---|---|---|---|---|---|---|---|---|---|---|---|---|---|---|---|---|
| | Citation | Time Unit | Model 1 | Model 2 | Model 3 | Model 1 | Model 2 | Model 3 | $k$ | $\rho$ | $\psi$ | $k$ | $\rho$ | $\mu$ | $\phi$ | $\chi$ | $\rho$ | | |
| Group 1 (see Figure 7) | | | | | | | | | | | | | | | | | | | |
| Acute Myeloid Leukemia (AML) | Ruggeri et al. | years | -1216 | -872 | -583 | -1203 | -859 | -570 | 1.433 | 0.53 | 0.0088 | 2.863 | 0.57 | 0.33 | 1.00 | 0.59 | 0.48 | 0.48 | 0.57 |
| Renal Cell Carcinoma | Marconi et al. | months | -1147 | -823 | -826 | -1134 | -810 | -813 | 0.023 | 0.57 | 0.0000 | 0.048 | 0.62 | 0.33 | 34.98 | 0.59 | 0.61 | 0.57 | 0.62 |
| Renal Cell Carcinoma | Zisman et al. | months | -1167 | -681 | -692 | -1154 | -668 | -679 | 0.037 | 0.33 | 0.0000 | 0.076 | 0.40 | 0.33 | 21.92 | 0.59 | 0.39 | 0.33 | 0.40 |
| Renal Cell Carcinoma | Brookman-Amissah et al. | months | -979 | -686 | -707 | -965 | -673 | -694 | 0.024 | 0.50 | 0.0000 | 0.053 | 0.56 | 0.33 | 29.68 | 0.59 | 0.56 | 0.50 | 0.56 |
| Hepatocellualr Carcinoma | Tran et al. | months | -1973 | -1725 | -1712 | -1960 | -1712 | -1699 | 0.041 | 0.87 | 1.9865 | 0.078 | 0.88 | 0.33 | 22.54 | 0.59 | 0.88 | 0.87 | 0.88 |
| Hepatocellualr Carcinoma | Sakon et al. | years | -1235 | -739 | -740 | -1222 | -726 | -727 | 0.346 | 0.16 | 0.0571 | 0.688 | 0.24 | 0.33 | 2.49 | 0.59 | 0.22 | 0.16 | 0.24 |
| Follicular Lymphoma | Kornacker et al. | months | -1309 | -991 | -1029 | -1296 | -978 | -1016 | 0.033 | 0.51 | 1.6284 | 0.062 | 0.56 | 0.33 | 27.70 | 0.59 | 0.54 | 0.51 | 0.56 |
| Basal Cell Carcinoma | Silverman et al. | years | -2030 | -1447 | -1423 | -2017 | -1434 | -1410 | 0.244 | 0.85 | 0.0001 | 0.545 | 0.87 | 0.33 | 3.18 | 0.59 | 0.87 | 0.85 | 0.87 |
| Non-small cell lung cancer | Maeda et al. | months | -1687 | -1493 | -1452 | -1674 | -1480 | -1439 | 0.013 | 0.75 | 0.8824 | 0.051 | 0.86 | 0.33 | 34.03 | 0.59 | 0.86 | 0.75 | 0.86 |
| Melanoma | Zalaudek et al. | months | -1889 | -1064 | -1782 | -1876 | -1051 | -1769 | 0.021 | 0.93 | 1.5043 | 99,948 | 0.97 | 0.17 | 40.85 | 0.59 | 0.94 | 0.93 | 0.97 |
| Melanoma | Morton et al. | years | -1504 | -1483 | -1432 | -1491 | -1470 | -1419 | 0.269 | 0.69 | 0.2898 | 0.498 | 0.72 | 0.33 | 3.57 | 0.59 | 0.71 | 0.69 | 0.72 |
| Chronic Myeloid Leukemia (CML) | Dulucq et al. | months | -609 | -680 | -688 | -596 | -667 | -675 | 0.190 | 0.47 | 1.4714 | 0.507 | 0.49 | 0.13 | 5.20 | 0.36 | 0.48 | 0.47 | 0.49 |
| Chronic Myeloid Leukemia (CML) | Ureshino et al. | months | -787 | -1201 | -1219 | -774 | -1188 | -1206 | 0.057 | 0.46 | 5.7260 | 0.174 | 0.55 | 0.10 | 16.25 | 0.31 | 0.55 | 0.46 | 0.55 |
| Chronic Myeloid Leukemia (CML) | Saussele et al. | months | -1002 | -1070 | -1152 | -989 | -1057 | -1139 | 0.347 | 0.52 | 0.5974 | 0.538 | 0.53 | 0.30 | 3.43 | 0.55 | 0.52 | 0.52 | 0.53 |
| Group 2 (see Suppl. Figure 1) | | | | | | | | | | | | | | | | | | | |
| Triple negative breast cancer | Bajpai et al. | months | -1258 | -1143 | -1152 | -1245 | -1130 | -1139 | 0.027 | 0.54 | 2.4254 | 0.052 | 0.59 | 0.33 | 34.31 | 0.59 | 0.57 | 0.54 | 0.59 |
| Triple negative breast cancer | Eom et al. | months | -1525 | -1473 | -1535 | -1512 | -1460 | -1522 | 0.033 | 0.80 | 5.4707 | 0.049 | 0.82 | 0.33 | 36.35 | 0.59 | 0.81 | 0.80 | 0.82 |
| Colorectal cancer | Liska et al. | years | -1819 | -1908 | -1807 | -1806 | -1895 | -1794 | 0.394 | 0.89 | 0.1364 | 0.796 | 0.90 | 0.33 | 2.25 | 0.59 | 0.90 | 0.89 | 0.90 |
| Non-small cell lung cancer | Xi et al. | months | -987 | -802 | -764 | -974 | -789 | -751 | 0.021 | 0.57 | 0.0000 | 0.045 | 0.64 | 0.33 | 38.30 | 0.59 | 0.62 | 0.57 | 0.64 |

| | | | | | | | | | | | | | | | | | | | |
|---|---|---|---|---|---|---|---|---|---|---|---|---|---|---|---|---|---|---|---|
| Non-small cell lung cancer (st. 1) | Taylor et al. | months | -1474 | -530 | -1485 | -1461 | -517 | -1472 | 0.004 | 0.00 | 6.2631 | 6438.0<br>50 | 0.89 | 0.25 | 66.13 | 0.59 | 0.66 | 0.00 | 0.89 |
| Non-small cell lung cancer (st. 2) | Taylor et al. | months | -1320 | -325 | -1162 | -1307 | -312 | -1149 | 0.008 | 0.00 | 3.9242 | 6437.1<br>10 | 0.81 | 0.24 | 48.07 | 0.59 | 0.55 | 0.00 | 0.81 |
