## Supplementary material for "The inherent fragility of collective proliferative control": Mathematical Appendix

# to

### Section 1. The variance of the distribution created by fine-tuned, balanced stem cell division grows without bound.

Here we seek to demonstrate that, when stem cells are independently dividing under homeostatic conditions, as long as they do not exhibit perfect division asymmetry (i.e. every stem cell generates exactly 1 stem cell in every division), then the variance in the number of stem cells grows without bound. In other words, if stem cells divide probabilistically, as long as they behave independently of each other, reliable homeostasis cannot be achieved.

We begin by considering a single stem cell at time $t$=0. Let its probability of producing two stem cells be $p_2$, its probability of producing one stem cell and one terminal cell be $p_1$ and its probability of producing two terminal cells be $p_0$, such that $p_0 + p_1 + p_2 = 1$. If $p_0 = p_2 = 0$, we have only asymmetric division, and the number of stem cells will stay constant through every division. Otherwise, stem cell numbers will evolve probabilistically, sometimes increasing and sometimes decreasing. If $p_0 = p_2$, then increases and decreases, should balance, on average, and it would seem the expectation value for the population size should remain constant. We seek, however, to determine how the distribution of population sizes evolves over time. Since $p_0 = p_2$, we will use $p$ to stand for both $p_0$ and $p_2$, and 1-2p for $p_1$.

We begin by understanding the time steps as corresponding to the division of a single stem cell, that is to say, every time a single stem cell divides, we increment $t$ by 1. Thus, with each step increment, the population of stem cells either grows by 1 (with probability $p$), shrinks by 1 (with probability $p$) or stays the same (with probability 1-2$p$). With respect to the distribution of probabilities of finding any given number of stem cells, it is obvious that steps in which the stem cell population neither grows nor shrinks have no effect on how that distribution evolves other than to introduce pauses, i.e., the time evolution will take longer but ultimately trace out the same path. Accordingly, if we are only concerned with the limit as $t$ approaches infinity, we may ignore such steps.

Knowing that asymmetric division has no effect on the limiting behavior enables us to simplify the problem by considering only the case in which $p$=1/2, i.e., no asymmetric division. We begin by writing down general equations for the evolution of the probability distribution of stem cells. Letting $\Pr[n,t]$ stand for the probability of finding $n$>0 stem cells at time $t$, we have:

$$\Pr[1,t] = p\Pr[2,t-1]$$
$$\Pr[n|n>1,t] = p\Pr[n-1,t-1] + p\Pr[n+1,t-1]$$

where $p$=1/2. Note that the first equation reflects the fact that the condition of $n$=1 can only arise by a decrease in stem cell number and not by an increase, since the state of $n$=0 (extinction) is an absorbing state.

Starting from initial conditions of $n$=1 at $t$=0, we enumerate below the probabilities of observing any $n$ for times up to $t$=12. In this matrix, the successive columns reflect time steps (from *t*=0 to *t*=12), and the rows are the probability of observing any number of stem cells (starting from 1 and increasing by 1 for each row) and *p*=1/2. We note that, for any *t*, the maximum possible *n* for which the probability is non-zero is *t*+1.

| | | | | | | | | | | | | |
|---|---|---|---|---|---|---|---|---|---|---|---|---|
| $1$ | $0$ | $p^2$ | $0$ | $2p^4$ | $0$ | $5p^6$ | $0$ | $14p^8$ | $0$ | $42p^{10}$ | $0$ | $132p^{12}$ |
| $0$ | $p$ | $0$ | $2p^3$ | $0$ | $5p^5$ | $0$ | $14p^7$ | $0$ | $42p^9$ | $0$ | $132p^{11}$ | $0$ |
| $0$ | $0$ | $p^2$ | $0$ | $3p^4$ | $0$ | $9p^6$ | $0$ | $28p^8$ | $0$ | $90p^{10}$ | $0$ | $297p^{12}$ |
| $0$ | $0$ | $0$ | $p^3$ | $0$ | $4p^5$ | $0$ | $14p^7$ | $0$ | $48p^9$ | $0$ | $165p^{11}$ | $0$ |
| $0$ | $0$ | $0$ | $0$ | $p^4$ | $0$ | $5p^6$ | $0$ | $20p^8$ | $0$ | $75p^{10}$ | $0$ | $275p^{12}$ |
| $0$ | $0$ | $0$ | $0$ | $0$ | $p^5$ | $0$ | $6p^7$ | $0$ | $27p^9$ | $0$ | $110p^{11}$ | $0$ |
| $0$ | $0$ | $0$ | $0$ | $0$ | $0$ | $p^6$ | $0$ | $7p^8$ | $0$ | $35p^{10}$ | $0$ | $154p^{12}$ |
| $0$ | $0$ | $0$ | $0$ | $0$ | $0$ | $0$ | $p^7$ | $0$ | $8p^9$ | $0$ | $44p^{11}$ | $0$ |
| $0$ | $0$ | $0$ | $0$ | $0$ | $0$ | $0$ | $0$ | $p^8$ | $0$ | $9p^{10}$ | $0$ | $54p^{12}$ |
| $0$ | $0$ | $0$ | $0$ | $0$ | $0$ | $0$ | $0$ | $0$ | $p^9$ | $0$ | $10p^{11}$ | $0$ |
| $0$ | $0$ | $0$ | $0$ | $0$ | $0$ | $0$ | $0$ | $0$ | $0$ | $p^{10}$ | $0$ | $11p^{12}$ |
| $0$ | $0$ | $0$ | $0$ | $0$ | $0$ | $0$ | $0$ | $0$ | $0$ | $0$ | $p^{11}$ | $0$ |
| $0$ | $0$ | $0$ | $0$ | $0$ | $0$ | $0$ | $0$ | $0$ | $0$ | $0$ | $0$ | $p^{12}$ |
| $0$ | $0$ | $0$ | $0$ | $0$ | $0$ | $0$ | $0$ | $0$ | $0$ | $0$ | $0$ | $0$ |
| $0$ | $0$ | $0$ | $0$ | $0$ | $0$ | $0$ | $0$ | $0$ | $0$ | $0$ | $0$ | $0$ |

The reason for the alternating zeros is that, by neglecting asymmetric divisions, each time step must produce either an increment of +1 or -1, so the distribution flips consecutively between having only odd-numbered values and only even-numbered ones. By inspection, we see that the nonzero coefficients may be expressed as

$\frac{n}{t+1}\binom{t+1}{\frac{t+1}{2}-\frac{n}{2}}p^t$ when *t* is odd, and $\frac{2n}{t+1+n}\binom{t}{\frac{t+1}{2}-\frac{n}{2}}p^t$ when *t* is even. It is straightforward to show that these forms satisfy the recursive equations for all *n*>0.

We now wish to show that, for *p*=1/2, the variance in the distributions defined by these expressions tends to infinity as *t* becomes large. Given a probability distribution function $P[t]$, the mean, aka the first moment, at time *t,* is simply $\sum_n nPr[n,t]$. The variance may then be found by subtracting the square of the mean from the second moment, which is $\sum_n n^2Pr[n,t]$.

First, we calculate the means for the two distributions in which *t* is either odd or even. In both cases, it is straightforward to show that the mean =1 independent of *t*. To do so, it is convenient to make some substitutions so that the iterators of summation increase in whole number steps, rather than in steps of size 2. For the case when *t* is odd, we define *d* = *n*/2 and $\tau$= (*t*+1)/2. Now the series *n*=2,4,6… becomes *d*=1,2,3…, and the series t=1,3,5… becomes $\tau$= 1,2,3… etc. The maximum *d* that produces a nonzero probability is then $\tau$.

Similarly, for the case when $t$ is even, we define $d = (n+1)/2$ and $\tau = 1+t/2$. Now the series $n$=1,3,5… becomes $d$=1,2,3…, and the series t=0,2,4… becomes $\tau$ = 1,2,3… etc. The maximum $d$ that produces a nonzero probability is then $\tau + 1$.

With these substitutions, and setting $p$=1/2, the sums that define the moments of these distributions are:

$$\sum_{d=1}^{\tau} \frac{2^{1+k-2\tau} d^{1+k}}{\tau} \binom{2\tau}{\tau - d}$$

when $t$ is odd and

$$\sum_{d=1}^{\tau+1} \frac{4^{-\tau}(2d-1)^{1+k}}{d+\tau} \binom{2\tau}{\tau - d + 1}$$

when $t$ is even, where $k$ is the moment number (i.e., $k$=1 returns the mean, and $k$=2 the second moment). With the assistance of *Mathematica* software, one may directly evaluate both expressions and show that, when $k$=1, both are equal to 1, for all $t \geq 0$.

When $k$=2, the expression for $t$-odd evaluates to a relatively simple form

$$2^{2-2\tau}(1+\tau)\binom{2\tau}{\tau-1}$$

the limit of which, as $\tau$ approaches ∞, is ∞. Thus, since the variance is the second moment minus the square of the mean, and since and the mean is a constant and $\tau$ is linearly related to $t$, the variance also approaches ∞ as $t \to \infty$.

When $k$=2, the expression for $t$-even evaluates to a more complicated form, the limit of which is difficult to calculate analytically. In this case it is simpler to show that the second moment when $t$ is even is always greater than the second moment when $t = t - 1$, implying that the second moment for even $t$ must also approach ∞ as $t \to \infty$.

To see this, consider any discrete probability distribution $P[n]$ with support on the nonnegative integers, and another one $Q[n]$ derived by averaging $P[n-1]$ and $P[n+1]$; we can show that the second moment of $Q[n]$ is always larger than the second moment of $P[n]$ and approaches ½ plus the second moment of $P[n]$ as $n$ becomes large. Specifically, if

$$Q[n] \equiv \frac{P[n-1] + P[n+1]}{2},$$

then the expression for the second moment of $Q$ may be rearranged as follows:

$$\sum_{n=1}^{\infty} n^2 Q[n] = \sum_{n=1}^{\infty} n^2 \frac{P[n-1]+P[n+1]}{2} = \frac{1}{2}\left(\sum_{n=1}^{\infty} n^2 P[n-1] + \sum_{n=1}^{\infty} n^2 P[n+1]\right)$$
$$= \frac{1}{2}\left(\sum_{n=0}^{\infty} (n+1)^2 P[n] + \sum_{n=2}^{\infty} (n-1)^2 P[n]\right)$$
$$= \frac{1}{2}\left(1^2 P[0] + \sum_{n=1}^{\infty} (n+1)^2 P[n] + \sum_{n=1}^{\infty} (n-1)^2 P[n] - 0^2 P[1]\right)$$
$$= \frac{1}{2}\left(P[0] + \sum_{n=1}^{\infty} n^2 P[n] + \sum_{n=1}^{\infty} 2nP[n] + \sum_{n=1}^{\infty} P[n] + \sum_{n=1}^{\infty} n^2 P[n] - \sum_{n=1}^{\infty} 2nP[n]\right.$$
$$\left. + \sum_{n=1}^{\infty} P[n]\right) = \frac{1}{2} P[0] + \sum_{n=1}^{\infty} n^2 P[n] + \sum_{n=1}^{\infty} P[n]$$

The term $P[0]$ is equivalent to $1 - \sum_{n=1}^{\infty} P[n]$. Thus, the above expression becomes

$$\frac{1}{2}\left(1 - \sum_{n=1}^{\infty} P[n]\right) + \sum_{n=1}^{\infty} n^2 P[n] + \sum_{n=1}^{\infty} P[n] =$$
$$\frac{1}{2} + \sum_{n=1}^{\infty} n^2 P[n] + \frac{1}{2}\sum_{n=1}^{\infty} P[n]$$

We note that the second of these three terms is exactly the second moment of $P[n]$. The third term is just a sum of non-negative numbers, so it is necessarily nonnegative. From this it follows that the second moment of $Q[n]$ is always greater than the second moment of $P[n]$, which is the required result.

One can also show that, for $p$=½, $\sum_{n=1}^{\infty} P[n]$ is equal to

$$\frac{4^{-\tau}(1+\tau)}{\tau}\binom{2\tau}{\tau-1}$$

which approaches zero as $\tau \to \infty$. This makes sense given that zero is an absorbing state, so that for $p \leq$ ½, all probability is eventually attracted there.

Putting this together indicates that, for sufficiently large *t*, the second moment of $Q[n]$ approaches ½ plus the second moment of $P[n]$. Thus, it is also the case for even values of *t* that variance grows without bound as t goes to infinity.

### Section 2. Feedback control of renewal by factors that diffuse in two-dimensional space is not globally stable.

Consider a growing disc with perfect cell mixing, in which renewal is controlled by negative feedback of terminal cells upon dividing cells, and where the feedback is mediated by a diffusible factor produced by terminal cells that rapidly diffuses and undergoes constant degradation or leaking. Due to the assumption of perfect mixing, the amount of feedback factor made at any point in a growing disc will be spatially uniform and proportional to the fraction of total cells in the disc that are terminal.

We know from Chen et al., (Chen et al., 2015) that the function describing the spatial gradient associated with uniform production within a disc is

$$v(1 - \rho\, I_0[x] K_1[\rho])$$

where $v$ is a rate that depends on what fraction of cells within the disc are producing the feedback factor (i.e. are terminal cells), $\rho$ is the radius of the disc, $x$ is a variable describing location along that radius, and $I_\alpha$ and $K_\alpha$ are Bessel functions of the first and second kind, respectively. In this formulation, $\rho$ and $x$ are expressed in spatial units equal to the decay length of the diffusible molecule.

It is clear that this expression is monotonic declining in $x$ because $I_0[x]$ is a monotonically increasing function in $x$ for $x$ >0.

Therefore, the minimum of this expression (i.e. the weakest feedback) will occur when $x = \rho$, i.e. at the edge of the disc. At that location, the value is

$$v(1 - \rho\, I_0[\rho] K_1[\rho])$$

Note that the expression $(1 - \rho\, I_0[\rho] K_1[\rho])$ is monotonically increasing in $\rho$, bounded between 0 (at $\rho$=0) and a limiting value of 1/2 as $\rho$ approaches ∞.

If the probability of renewal at that location, which we will call $\mathcal{P}[\rho]$, is a Hill function of the feedback, we may write:

$$\mathcal{P}[\rho] = \frac{p_0}{1 + gv(1 - \rho I_0[\rho] K_1[\rho]))}$$

Where $p_0$ is a constant between 0.5 and 1, and $g$ is a nonnegative constant representing the feedback strength. The minimum value of $\mathcal{P}[\rho]$ will clearly occur when $v$ is maximal, i.e., when all of the cells at or near the boundary ($x = \rho$) are terminal. We shall call that value of $v$, $v_{\max}$. So, the minimum possible value of $\mathcal{P}[\rho]$ is thus:

$$\frac{p_0}{1 + gv_{\max}(1 - \rho I_0[\rho] K_1[\rho])}$$

Now we will ask whether there is always a value of g for which this expression must necessarily be greater than 1/2, independent of $\rho$. If so, then cells at the boundary will necessarily grow without bound. We see by inspection that this expression will be greater than 1/2 provided

$$g < \frac{2p_0 - 1}{(1 - \rho I_0[\rho]K_1[\rho])v_{\max}}$$

We also note that the maximum possible value of $1 - zI_0[z]K_1[z]$ is 1/2. Therefore choosing

$$g < \frac{2\mathrm{p}0 - 1}{(1/2)v_{\max}}$$

guarantees that, regardless of $\rho$, $\mathcal{P}[\rho]$ will necessarily be greater than 1/2. Since $2p_0 - 1$ and $v_{\max}$ are both positive numbers, it will always be possible to find a non-zero $g$ that satisfies this condition. In other words, if feedback strength is weak enough, neither a steady state nor final state can be reached. *QED*.

A similar set of calculations can be made for the three-dimensional scenario, in which case the equation for the level of diffusible molecule at the boundary is

$$2\nu(1 - \frac{(1+\rho)}{\rho(1 + \mathrm{Coth}[\rho])})$$

### Section 3. Growth control in an epithelium modeled as a flat sheet of semi-infinite extent

As an alternative to modeling a tissue as a growing ball, we can use a flat geometry in which all dividing cells reside close to a basement membrane, to mimic a typical epithelium. To model this three dimensional arrangement using differential equations it is necessary to specify six boundary conditions, indicating what happens to diffusing substances at the apical surface, the basal surface, and in the four planar directions along the length and width of the epithelium. If the length and width of the epithelium are much larger than the characteristic decay length of any diffusing substances (defined as the square root of the diffusivity divided by the substance's rate of decay), then outcomes throughout most of the epithelium will be independent of what happens at the planar boundaries, and such boundaries may be ignored. This transforms the problem into a one-dimensional one (along the apico-basal dimension), thus requiring only two boundary conditions. As epithelia are often centimers or longer in planar dimension, and diffusing biomolecules tend to have decay lengths of at most hundreds of microns, this one-dimensional approximation is likely a good one for most epithelia.

At the apical surface of an epithelium it is natural to assume a no-flux boundary condition, as apical surfaces are commonly sealed by tight junctions that stop the spread of intercellular molecules. At the basal surface it is natural to assume that there is no obstacle to diffusion, as the porosity of basement membranes typically allows free diffusion of protein-sized molecules. Mathematically, a "non-boundary" boundary like this is handled by placing a fictitious boundary far away, solving one's equations, and then taking the limit as that boundary goes to infinity.

If we use *x* to represent position (with *x* =0 corresponding to the basal surface), *h* to represent the total height of the epithelium (i.e. the apicobasal distance) and $\alpha$ to represent the height of the dividing cell layer that lies next to the basement membrane, then we may derive the following equation for the steady state gradient associated with any diffusible substance produced by the non-dividing cell layer.

$$\nu e^{\frac{x-\alpha}{\lambda}}\left(\frac{1}{1+\coth[\frac{h-\alpha}{\lambda}]}\right) \qquad 0 \le x < \alpha$$

$$\nu\left(1-\frac{\cosh\left[\frac{h-x}{\lambda}\right]\operatorname{sech}\left[\frac{h-\alpha}{\lambda}\right]}{1+\tanh\left[\frac{h-\alpha}{\lambda}\right]}\right) \quad \alpha \le x \le h$$

where $\nu$ is the rate of production and $\lambda$ is the characteristic decay length of the substance (which we assume to be constant everywhere). This is obtained by solving the pair of diffusion equations $0 = \nu - c[x] + \lambda^2 c''[x], 0 \le x < \alpha,$ and $0 = -c[x] + \lambda^2 c''[x], \alpha \le x < h$ and requiring agreement of both value and derivative at $x$=$\alpha$. Since we only need to concern ourselves with the values in the range $0 \le x < \alpha$ (where the dividing cells are), the only expression needed is the first of those above. Assuming this represents the concentration of an inhibitory factor, then the local value of the renewal probability, at any location x, may be modeled as:

$$p[x] = \frac{\bar{p}}{1+\frac{e^{\frac{x-\alpha}{\lambda}}\gamma}{1+\coth[\frac{h-\alpha}{\lambda}]}}$$

Where $\bar{p}$ is the maximum possible value of the renewal probability and g is a constant representing the product of $\nu$ and the strength of the negative feedback. We may find the average of this by integrating it from 0 to $\alpha$ and dividing by $\alpha$.

$$\bar{p}\frac{\lambda}{\alpha}\ln\left[\frac{\gamma + e^{\alpha/\lambda}(1+\coth[\frac{h-\alpha}{\lambda}])}{\gamma+1+\coth[\frac{h-\alpha}{\lambda}]}\right]$$

If we wish to represent this expression using numbers of cells, rather than measures of epithelial thicknesses ($h$ and $\alpha$), we note that $\alpha$ and h-$\alpha$ are just equal to c0 and c1, respectively, times a proportionality constant. The proportionality constant is simply a measure of how much planar area of the epithelium is being considered when enumerating cells. For example, if we are considering an area of epithelium = w, and it's height is $\alpha$, then the number of cells in it is w*$\alpha$ /$V_{cell}$, where V is the volume of a single cell. Thus we may replace $\alpha$ with c0 $V_{cell}$ /w and h-$\alpha$ with c1 $V_{cell}$ /w. If we define $\kappa$ as the unitless ratio $\lambda$ w/ $V_{cell}$ , we may re-write the above expression as

$$\bar{p}\frac{\kappa}{c0}\ln\left[\frac{\gamma + e^{c0/\kappa}(1+\coth[\frac{c1}{\kappa}])}{\gamma+1+\coth[\frac{c1}{\kappa}]}\right]$$

The dynamics of the entire system are therefore described by:

$$c0'[t] = c0\left(\frac{2\bar{p}\kappa}{c0}ln\left[\frac{\gamma + e^{c0/\kappa}(1+\coth[\frac{c1}{\kappa}])}{1+\gamma+\coth[\frac{c1}{\kappa}]}\right]-1\right)$$

$$c1'[t] = 2*c0\left(1-\frac{\bar{p}\kappa}{c0}ln\left[\frac{\gamma + e^{c0/\kappa}(1+\coth[\frac{c1}{\kappa}])}{1+\gamma+\coth[\frac{c1}{\kappa}]}\right]\right)-d*c1$$

where $d$ is the rate constant for turnover of the differentiated (terminal) cells. Note that if we assume cells are roughly spherical with radius r, $V_{cell} = \frac{4\pi r^3}{3}$, and if the area of epithelium being considered is circular with radius R, then $\kappa = \frac{3}{4}\left(\frac{R}{r}\right)^2\frac{\lambda}{r}$. So if $\lambda$ is 20 cell radii, and the region of epithelium being considered is a circle with a radius of 10 cell radii, then $\kappa$ = 1500. So it is reasonable to use values of $\kappa$ that are large (e.g., greater than 100).

### Section 4. Growth control in three-dimensional spherical geometry

As in Figures 2 and 4, we can use ordinary differential equations to model spatial conditions in which dividing and differentiated cells either fully mix, or fully sort away from each other. In Fig. 2 and 4 we consider a two-dimension disc geometry; here we approach the same problem in three dimensions. In both cases, the equations governing how negative feedback affects the renewal probability of dividing cells are the same, based on the idea that differentiated cells release a diffusible factor that is subject to constant decay over time.

The difference lies in how the average renewal probability gets calculated. In spherical geometry, the function describing the steady state diffusion gradient inside and outside a producing sphere is (Chen et al., 2015): $2\nu - \frac{2\nu(1+b\rho)\mathrm{csch}[b]\sinh[R]}{R(\rho+\coth[b])}$ inside the sphere and $\frac{2e^{(b-R)\rho}\nu(-1+b\coth[b])}{R(\rho+\coth[b])}$ outside the sphere, where

$\nu$ = rate of production

b = $\beta/\lambda_{\text{in}}$ where $\beta$ = the radius of the sphere

$\rho = \lambda_{\text{in}} / \lambda_{\text{out}}$

R = $r / \lambda_{\text{in}}$, where $r$ = radial position

To find the average value of a function $f$[r] in spherical coordinates, you need to calculate the triple integral of the function over the desired spherical region, then divide by the volume of that region; for a sphere of radius $\beta$, and a function that's spherically symmetrical, this becomes

$$< f >= \int_0^{\beta} 4\pi\rho^2 f[\rho]\, d\rho$$

Let A be the number of dividing cells, and B be the number of differentiated cells. For convenience we will say that each cell has radius 5. Then the volume for "$n$" cells is n * $4\pi/3$ * $5^3$. If all the cells are together in one sphere, then the radius of that entire sphere is $5n^{1/3}$

Let k stand for the inverse of the feedback strength. Thus, the equation for feedback would be:

$$\frac{kp}{k + F[R]}$$

Where $p$ stands for the maximum renewal probability.

<u>Well-mixed case:</u> A and B cells are thoroughly mixed. B cells produce feedback at a constant rate, but the feedback at any location in space reflects the proportion of cells that are B, as opposed to A (i.e. B/(A+B)).

The equation for F uses the inner solution which is $2\nu - \frac{2\nu(1+b\rho)\mathrm{Csch}[b]\mathrm{Sinh}[R]}{R(\rho+\mathrm{Coth}[b])}$.

For simplicity, we will take $\rho$ = 1. Replacing b with $\beta/\lambda$ and R with $r/\lambda$, we get F =

$$2\nu\left(1 - \frac{(1+\frac{\beta}{\lambda})\mathrm{csch}[\frac{\beta}{\lambda}]\sinh[\frac{r}{\lambda}]}{\frac{r}{\lambda}(1+\coth[\frac{\beta}{\lambda}])}\right)$$

Into which $\nu$ is replaced by B/(A+B) and $\beta$ is replaced by $5(A+B)^{1/3}$. Accordingly, the average value of the renewal probability experienced by A cells is:

$$\frac{3p}{5^3(A+B)}\int_0^{5(A+B)^{1/3}} \frac{r^2 * k}{k + (\frac{B}{A+B})\left(1 - \frac{(1+\frac{5(A+B)^{1/3}}{\lambda})\mathrm{csch}[\frac{5(A+B)^{1/3}}{\lambda}]\sinh[\frac{r}{\lambda}]}{\frac{r}{\lambda}(1+\coth[\frac{5(A+B)^{1/3}}{\lambda}])}\right)}dr$$

<u>Differentiated cells sort to the inside:</u>

The equation for F is now the outer solution. We make the same substitutions as before, but $\nu$=1, because in the producing region there are only differentiated cells. Accounting for the volume of the producing region being $5B^{1/3}$, we must integrate from $5B^{1/3}$ to $5(A+B)^{1/3}$, and the volume of the region of integration is $\frac{4}{3}\pi(5^3A)$. Altogether, the average renewal probability becomes:

$$\frac{3p}{5^3(A)}\int_{5B^{1/3}}^{5(A+B)^{1/3}} \frac{r^2 * k}{k + \left(\frac{2e^{(\frac{5(B)^{1/3}}{\lambda}-\frac{r}{\lambda})}(-1+\frac{5(B)^{1/3}}{\lambda}\coth[\frac{5(B)^{1/3}}{\lambda}])}{\frac{r}{\lambda}(1+\coth[\frac{5(B)^{1/3}}{\lambda}])}\right)}dr$$

<u>Differentiated cells sort to the outside:</u>

The equation for F is now based on the inner solution. To calculate the amount of production produced by an annular shell, we subtract the result for the entire sphere from the result for the region where dividing cells are. We make the same substitutions as before, with $\nu$=1, and the volume of the region of integration is again $\frac{4}{3}\pi(5^3A)$. The limits of integration are from zero to $5A^{1/3}$, so that altogether the average renewal probability becomes

$$\frac{3p}{5^3(A)}\int_0^{5A^{1/3}} \frac{r^2 * k}{k + \left(1 - \frac{(1+\frac{5(A+B)^{1/3}}{\lambda})\mathrm{csch}[\frac{5(A+B)^{1/3}}{\lambda}]\sinh[\frac{r}{\lambda}]}{\frac{r}{\lambda}(1+\coth[\frac{5(A+B)^{1/3}}{\lambda}])}\right) - \left(1 - \frac{(1+\frac{5(A)^{1/3}}{\lambda})\mathrm{csch}[\frac{5(A)^{1/3}}{\lambda}]\sinh[\frac{r}{\lambda}]}{\frac{r}{\lambda}(1+\coth[\frac{5(A)^{1/3}}{\lambda}])}\right)}dr$$

### Section 5. The impact of stem cell self-regulation on tissue growth control

Allowing terminal cells to negatively feed back onto the renewal probabilities of dividing cells ("renewal control") is not the only type of feedback that could occur in stem/terminal cell systems. Indeed, with additional complexity—such as multi-stage or branched lineages—many possible types of regulation could be envisioned. Several of these have been discussed in other studies, and in several cases the long-term behaviors of steady-state and final-state systems have been calculated (e.g. Buzi et al., 2015; Kunche et al., 2016; Lo et al., 2009). Here we consider the possibility that stem cells receive some type of negative feedback from themselves, and not just from terminal cells. For simplicity, we shall limit discussion to steady state systems (i.e., systems in which terminal cells turn over).

An important finding is that not all forms of regulation are capable of producing a steady state, and of those, not all produce a robust steady state. By robust, we mean a steady state in which small changes to the cell cycle speed of stem cells, or the turnover rate of terminal cells, do not produce significant changes to the final number of terminal cells. *In vivo* observations support the view that numbers of differentiated cells of tissues and organs are (and need to be) robust in this sense, because external factors (injury, viral infection, temperature, hormonal fluctuation, etc.) commonly influence cell turnover and/or cell division rates in unpredictable ways, yet tissues sizes are generally well maintained.

This type of robustness is best quantified using the Sensitivity Coefficient, $S$, a measure of the fold change in a response given a small fold change in a parameter. $S = 1$ means the response varies linearly (and directly) with the parameter whereas $S = -1$ means the response varies linearly (and inversely) with the parameter. The closer the value of $S$ to zero, the more robust the system.

<u>Renewal control (regulation of renewal probability by terminal cells)</u>
We begin by reviewing the findings with renewal control, which in ODE form may be modeled as

$$\mathrm{c0}'[t] = v\mathrm{c0}\left(-1 + \frac{2p}{1+\gamma \mathrm{c1}}\right),\quad \mathrm{c1}'[t] = 2v\mathrm{c0}\left(1 - \frac{p}{1+\gamma \mathrm{c1}}\right) - d\mathrm{c1}$$

Where $v$ is the rate of the cell cycle, *d* is the rate of terminal cell death, $\gamma$ is the strength of feedback and *p* the maximum renewal probability. The steady state solution for $\mathrm{c1}$ is $\frac{2p-1}{\gamma}$. The fact that neither $v$ nor $d$ occur in that formula means that terminal cell numbers are perfectly robust to both the cell cycle speed ($v$) and the rate of terminal cell turnover, $d$. Perfect robust adaptation reflects the fact that renewal control implements integral negative feedback (Buzi et al., 2015; Gupta and Khammash, 2022).

<u>Stem cell self-regulation of renewal probability</u>
Here the only feedback comes from stem cells, but it acts by lowering the renewal probability rather than the cell cycle speed. We model this scenario with the equations:

$$\mathrm{c0}'[t] = v\mathrm{c0}\left(-1 + \frac{2p}{1+\gamma \mathrm{c0}}\right),\ \mathrm{c1}'[t] = 2v\mathrm{c0}\left(1 - \frac{p}{1+\gamma \mathrm{c0}}\right) - d\mathrm{c1}$$

This system has a steady state in which $\mathrm{c1}$ is $\frac{(2\mathrm{p}-1)v}{d\gamma}$. As this varies linearly with both $v$ and $d$, i.e. $S = 1$ or $S = -1$, respectively, the steady state is non-robust and consequently not suitable for producing robust homeostasis.

<u>Stem cell self-regulation of cell cycle speed</u>

Here the only feedback comes from stem cells, which decrease their own rate of division as their numbers grow. We model this scenario with the equations:

$$\mathrm{c0}'[t] = \frac{(-1+2p)\mathrm{c0}\bar{v}}{1+\eta\mathrm{c0}}, \mathrm{c1}'[t] = \frac{2(1-p)\mathrm{c0}\bar{v}}{1+\eta\mathrm{c0}} - d\mathrm{c1}$$

Where $\bar{v}$ is the maximum possible cell cycle speed, and $\eta$ is the feedback strength. These equations do not produce a steady state, but rather one in which stem cell numbers rise without bound. On its own, therefore, this strategy does not produce control.

<u>Stem cell self-regulation of both renewal probability and cell cycle speed</u>
Here we combine both of the above types of feedback. Now the equations become:

$$\mathrm{c0}'[t] = \frac{\bar{v}}{1+\eta\mathrm{c0}}\frac{(-1+2p)\mathrm{c0}}{1+\gamma\mathrm{c0}}, \mathrm{c1}'[t] = \frac{\bar{v}}{1+\eta\mathrm{c0}}\frac{2(1-p)\mathrm{c0}\bar{v}}{1+\gamma\mathrm{c0}} - d\mathrm{c1}$$

These equations do produce a steady state, but it is just as non-robust to $\bar{v}$ and $d$ as in the case where stem cells feedback only on the renewal probability.

This analysis suggests that stem cell self-regulation, on its own, is not a viable strategy for producing robust, steady-state tissues. Instead, some sort of feedback regulation by terminal cells seems to be required. Nevertheless, mixing stem cell self-regulation together with renewal control from terminal cells might have desirable properties in terms of influencing system dynamics, for example altering the probability of stochastic escape from control. We therefore consider next such mixed strategies.

<u>Combining stem cell self-regulation of renewal probability with renewal control from terminal cells</u>
Here feedback from stem cells and feedback from terminal cells both lower the renewal probability. We model this scenario with the equations:

$$\mathrm{c0}'[t] = v\mathrm{c0}(-1+\frac{2p}{1+\eta\mathrm{c0}+\gamma\mathrm{c1}}),\ \ \mathrm{c1}'[t] = 2v\mathrm{c0}(1-\frac{p}{1+\eta\mathrm{c0}+\gamma\mathrm{c1}}) - d\mathrm{c1}$$

Here we find that the steady state solution for $\mathrm{c1}$ is $(\frac{v(-1+2p)}{v\gamma+d\eta})$, while that for $\mathrm{c0}$ is $(\frac{d(-1+2p)}{v\gamma+d\eta})$, so both have an explicit dependence on $v$ and $d$. We may lump $v$ and $d$ together into a single parameter $\delta = d/v$, and also define a parameter $\alpha = \eta/\gamma$. We find that the Sensitivity coefficients for the sensitivity of $\mathrm{c0}$ and $\mathrm{c1}$ to $\delta$ are $\frac{1}{1+\alpha\delta}$ and $-\frac{\alpha\delta}{1+\alpha\delta}$, respectively. We plot these below for three different values of $\alpha$.

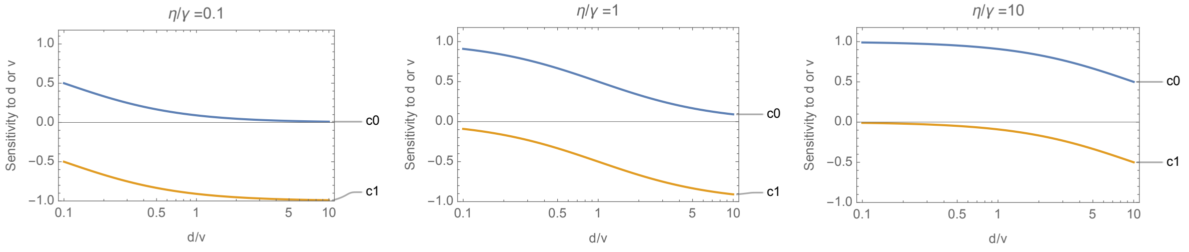

We note at once that it is impossible for both c0 and c1 to be robust, as the sum of the absolute values of their sensitivity coefficients always equals 1. Moreover, to achieve good robustness of c1, it is necessary for $\eta/\gamma \geq 1$, which means at least half of the feedback must come from terminal cells. Under these circumstances, the effect of feedback on the dynamics of approach to steady state is relatively small. For example, suppose we define "robust to c1" as meaning a sensitivity coefficient whose absolute value is less than 0.15. Comparing a scenario without feedback from stem cells, to a scenario with such feedback, and constraining them both to achieve the same steady state value of c1, as well as a sensitivity coefficient for c1 less than or equal to 0.15, sets a maximum on the value of $\eta$.

Specifically, if the case without feedback from stem cells uses parameters $\gamma 1$ for $\gamma$ and 0 for $\eta$, while the case with feedback from stem cells uses $\gamma 2$ for $\gamma$ and $\eta 2$ for $\eta$, we may solve for the values of $\gamma 2$ and $\eta 2$ that achieve both the desired steady state and the desired level of robustness. An example is shown below.

Here, $v = 1, d = 0.2, p = 0.9, \gamma = 0.005, \eta = 0$. In the case at right, $\eta = 0.00375$, the highest value consistent with adequate robustness of c1, and $\gamma$ is adjusted to 0.00425, which preserves the steady state value of c1. There is a subtle change to the dynamics of approach to steady state.

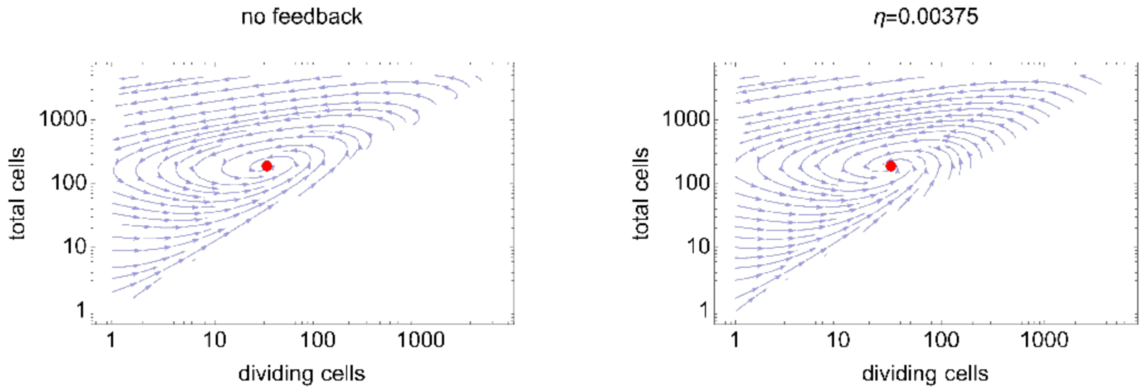

We can exaggerate the effect by increasing $\eta$ but now c1 is no longer robust. For example, in the case below with stem cell feedback $\eta = 0.02$ and $\gamma = 0.001$, we find the streamlines head more directly to the steady state, i.e. they oscillate less, which we might expect to reduce the magnitude of stochastic fluctuations away from steady state. The price for this, however, is a strong loss of robustness. For example, for these parameters the absolute value of the sensitivity of c1 to *d* or *v* is 0.8, meaning a two-fold increase in *d* would cause a 74% reduction in the steady state value of c1.

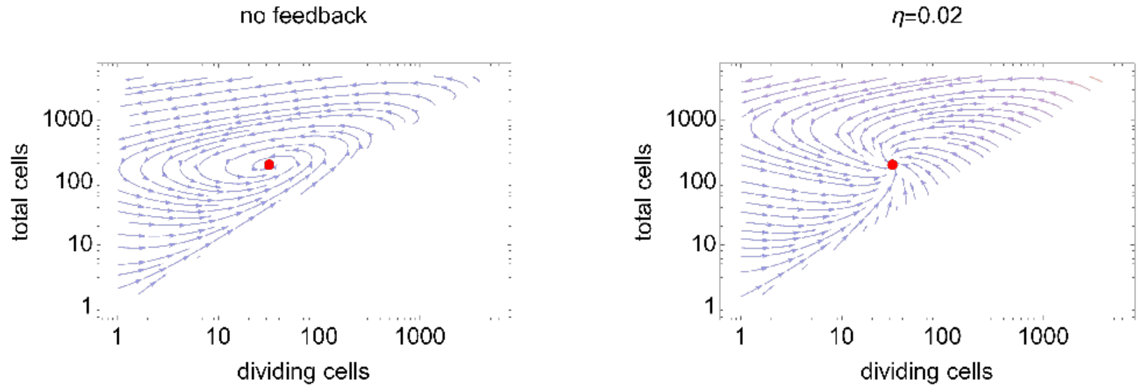

How might such behavior influence stochastic escape from control? To model this, we turn to the scenario considered in Figure 5, where terminal cells provide both positive and negative feedback, creating a bifurcation that allows from stochastic escape. In particular, we consider the situation modeled in Figure 5G. We next ask what happens when we add negative feedback from stem cells on the renewal probability. This generates the following equations:

$$c0'[t] = vc0\left(-1 + \frac{2p}{1 + \frac{\eta c0 + \gamma c1}{1 + (\varphi c1)^h}}\right), c1'[t] = 2vc0\left(1 - \frac{p}{1 + \frac{\eta c0 + \gamma c1}{1 + (\varphi c1)^h}}\right) - dc1$$

where, as before, $\eta$ captures the strength of stem cell feedback. The left panel below shows the stream plots that describe the behavior of this system when there is no stem cell feedback ($\eta = 0$). In the right panel, $\eta$ and $\gamma$ were adjusted to produce the identical steady state number of c1 cells, while using the maximum value of $\eta$ compatible with a sensitivity coefficient of 0.15 (for the absolute value of the sensitivity of c1 to $v$ and $d$). Notice that a separatrix appears in both cases. The parameter values used for these simulations were $v = 1, d = 0.1, p = 0.9, h = 1.5, \varphi = 0.0012, \gamma = 0.00187, \eta = 0$ (left) and $v = 1, d = 0.1, p = 0.9, h = 1.5, \varphi = 0.0012, \gamma = 0.00182, \eta = 0.0005$ (right)

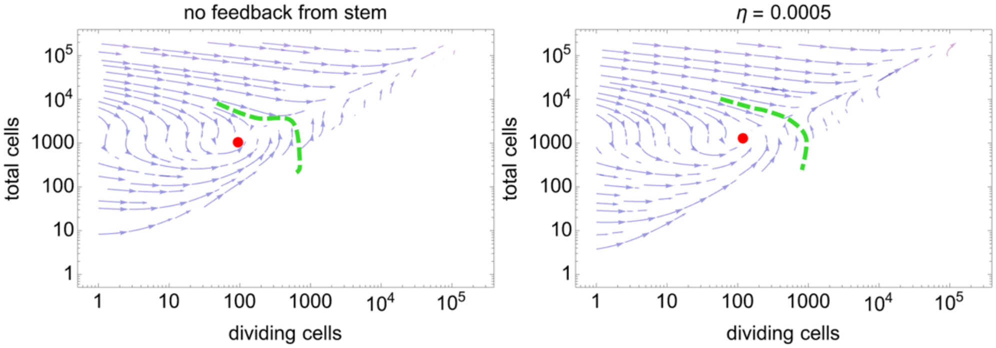

One way to examine the impact of the small changes to dynamics implied by these stream plots on the probability of stochastic escape from growth control is to initialize stochastic simulations from the calculated steady state values and run them for sufficiently long times to observe a substantial number of escapes (as in Figures 6A and 7A). Below this was done for the two cases shown above, simulating 100 runs of 1000 cell cycles each in both cases. In the case with no feedback from stem cells, 15 escape events were observed. With stem cell feedback, 8 such events were observed. Thus, stem cell feedback on renewal can modestly lower the probability of escape.

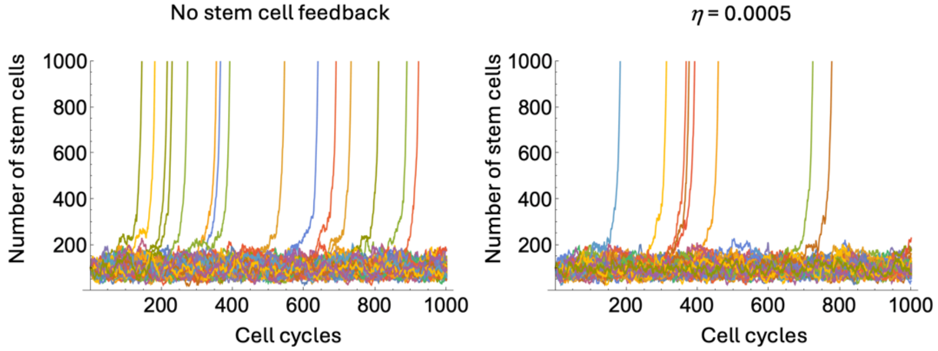

Combining stem cell self-regulation of cell cycle speed with renewal control from terminal cells

Lastly, we consider what happens when feedback from stem cells lowers the cell cycle speed, and feedback from terminal cells lowers the renewal probability. We model this scenario with the equations:

$$c0'[t] = c0\left(-1 + \frac{2p}{1 + \gamma c1}\right)\left(\frac{\bar{v}}{1 + \eta c0}\right),$$

$$c1'[t] = 2c0(1 - \frac{p}{1 + \gamma c1})(\frac{\bar{v}}{1 + \eta c0}) - dc1$$

In this case, the steady state number of terminal cells, c1, is $\frac{2p-1}{\gamma}$, the same as in the case of simple renewal control, so c1 is perfectly robust. The main thing that stem cell self-regulation does is slow the dynamics of the approach to steady state. This reduces oscillations about the steady state, but with increasing strength of feedback, an ever larger number of stem cells is required to support the same number of terminal cells, until eventually, with strong enough feedback, no number of stem cells will suffice, and no steady state can be reached. We may visualize this process by examining stream plots analogous to those in Figure 2, with increasing values of $\eta$, the strength of stem cell feedback. The red dot shows the position of the steady state, which vanishes when $\eta$=0.07. In these plots, $\bar{v} = 1, d = 0.2, p = 0.9,$ and $\gamma = 0.01$.

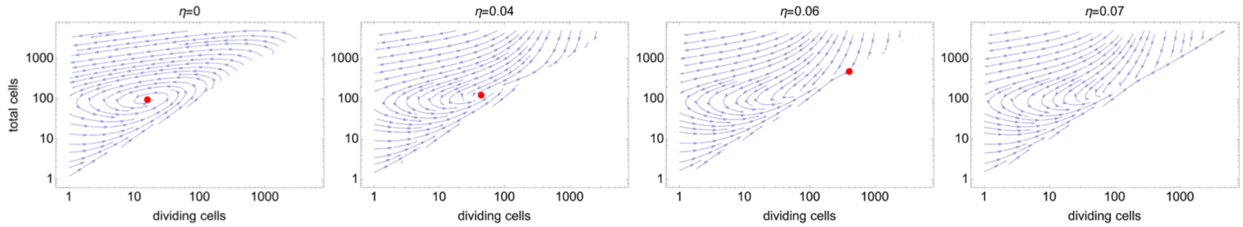

Thus, stem cell feedback of this type doesn't improve the robustness of the steady state, delays acquisition of the steady state, requires a large number of stem cells to support the generation of terminal cells, and creates parameter regimes that are unstable.

Next, we may ask what influence stem cell feedback of this type would have on the probability of stochastic escape from growth control. To do so, we once again consider a situation where there is explicit positive feedback, of the type in Figure 5G. Specifically, we allow mixed negative and positive feedback from terminal cells and ask what happens when we add negative feedback from stem cells on the rate of proliferation, $\bar{v}$. We model this scenario with the equations:

$$c0'[t] = \frac{\bar{v}}{1+\eta c0} c0 \left( -1 + \frac{2p}{1 + \frac{\gamma c1}{1+(\varphi c1)^h}} \right),$$

$$c1'[t] = 2\frac{\bar{v}}{1+\eta c0} c0[t] \left( 1 - \frac{p}{1 + \frac{\gamma c1}{1+(\varphi c1)^h}} \right) - d c1$$

As before, $\gamma$ is the strength of negative feedback from terminal cells, $\varphi$ is the strength of positive feedback from terminal cells, and $\eta$ is the strength of feedback from stem cells on the rate of proliferation. As previously, the steady state solution for c1 is independent of $\eta$, i.e., feedback from stem cells only changes the steady state number of stem cells. When we examine the dynamics of escape, we see that the probability of escape—as estimated from stochastic simulations like those in Fig. 5—declines with increasing $\eta$.

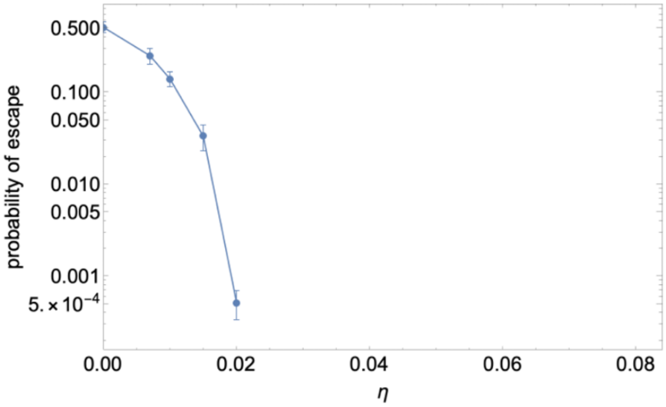

In this particular example, the parameters were $v = 1, d = 0.2, p = 0.9, \gamma = 0.018, \varphi = 0.0098, h = 2$. The abscissa on this plot extends to $\eta \approx 0.0839$, because that is the point at which the system becomes completely unstable (to achieve a steady state the fraction of stem cells needs to exceed 100%, which is impossible). This raises the question of whether, between the value of $\eta \approx 0.02$ and $\eta \approx 0.0839$, stem cell feedback can drive the probability of escape to zero, or just a very small number. Although we cannot address this question analytically for arbitrary values of $h$, for $h = 2$ we can do so. In this case, directly solving the system equations with rates set to zero returns three steady states: a trivial stable state where c0=0 and c1=0, a stable steady state, and an unstable steady state. As long as the unstable steady state exists there will be a separatrix that divides the systems into regimes of stability and uncontrolled growth. However, for $h = 2$ it is straightforward to show that the unstable steady state vanishes for

$$\eta > \frac{v}{2d}\left(\frac{\gamma}{(2p-1)} - \sqrt{\left(\frac{\gamma}{2p-1}\right)^2 - 4\varphi^2}\right)$$

Whereas obligatory unbounded growth does not occur until

$$\eta > \frac{v}{2d}\left(\frac{\gamma}{(2p-1)} + \sqrt{\left(\frac{\gamma}{2p-1}\right)^2 - 4\varphi^2}\right)$$

In effect, the system can take on three possible configurations, depending on parameters. For sufficiently small $\eta$ it is bi-modal, exhibiting either stability or unbounded growth depending on initial conditions; for somewhat larger $\eta$ it can be monostable provided $\gamma$ is not too small, and for still larger $\eta$ it is categorically unstable, always exhibiting unbounded growth.

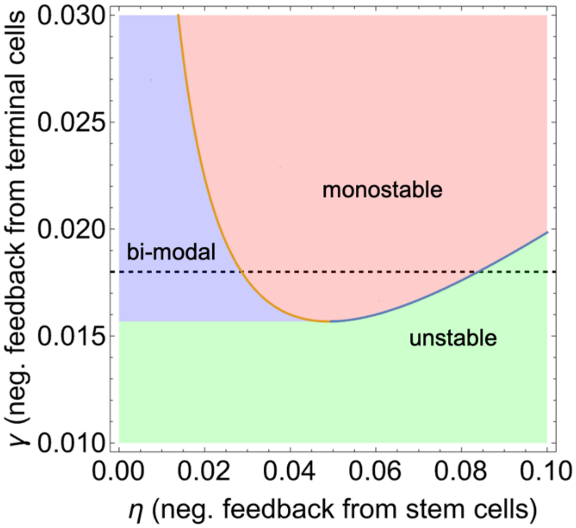

The above bifurcation diagram shows the locations of these three regions with respect to the parameters $\eta$ and $\gamma$, for the conditions $v = 1, d = 0.2, p = 0.9, \varphi = 0.0098$. The dashed line shows the results when $\gamma = 0.018$, as in the scenario plotted above.

From this analysis, we conclude that feedback of stem cell numbers on their own rate of proliferation can be an effective tool for reducing the probability of escape from control, but only at the expense of the creation of a zone of instability, even when $\gamma$ is large enough to otherwise preclude such instability.

Because this is a simplified model, it is difficult to know how effective stem cell feedback could be in real world situations, as the model assumes stem cell feedback rises linearly with the number of stem cells, which would be unlikely to be the case when signals must spread through space. For example, if feedback is mediated by direct contact between stem cells—as might be the case with Notch signaling, or signaling through protocadherins of the Fat family—one would expect feedback to saturate at rather low numbers of stem cells, hence it would rise much slower than linearly under many conditions. One might think this could be compensated for by increasing the strength of feedback but, as shown above, negative feedback from stem cells that is too strong will lead to instability on its own.

In the end it is interesting to speculate that mutation of genes involved in stem cell self regulation, such as Notch, might move a system from the monostable to the bimodal regime of the above bifurcation diagram. The prediction is that this would have little phenotypic effect (as the numbers of c1 cells, the predominant cell type, are independent of $\eta$), but might set the system up for later, stochastic escape. This could explain why, for example, oncogenic *NOTCH* mutations are frequently found in phenotypically normal skin (e.g., Martincorena et al., 2015).

### Section 6. Models of tumor re-growth

For each potential mechanism of recurrence, we develop a model of the survival function, *f*[$t$], the probability that a treated patient is still tumor-free after any given amount of time. We begin by allowing for the possibility that some fraction of patients, $\rho$, has been "cured", i.e. treatment has completely removed all cancer cells and therefore only the remaining fraction (1- $\rho$), which we will refer to as the susceptible fraction, has a risk of recurrence.

**Model 1: Stochastic emergence from dormancy**

We consider here that case in which, following therapy, any tumor residuum is dormant and has a constant probability of starting to grow again, after which it simply grows exponentially. As the waiting time distribution for a constant probability event is the exponential distribution, the probability that a tumor has initiated by time $t$ is simply the CDF of the exponential distribution, i.e., $1 - e^{-kt}$, where $1/k$ is the average waiting time between events. Thus, the survival function for tumor recurrence in the susceptible pool is $e^{-kt}$. However, we must account for the fact that, clinically, we do not detect tumors at the moment they recur, but only after they have regrown to exceed some threshold of detectability. If that threshold size is $\phi$ times the size at which initiation occurs, then the time delay from initiation (e.g. 1 cell) to detection $\phi$ cells) is just the time it takes to grow $\phi$-fold, which we shall call $\psi$. Thus, the survival function for susceptible patients will be $e^{-k(t-\psi)}, t > \psi$. To fit data that include both susceptible and non-susceptible patients, the full expression should be $\rho + (1-\rho)e^{-k(t-\psi)}$.

**Model 2: Variable tumor residuum size**

Here we consider that all tumors grow at about the same exponential rate, characterized by rate constant $k$, and therapy removes some fraction of the tumor, leaving behind a residuum that simply regrows and is detected once it reaches detectable size. Again, we will consider the possibility that a fraction $\rho$ of patients is cured, *i.e.,* had no tumor residuum, so that re-growth applies only to the susceptible fraction.

Assuming a residuum size $s$, the time it takes to grow to any threshold size $\theta$ is just $\frac{1}{k} ln \frac{\theta}{s}$. If we define $m=s/\theta$, i.e., scale residuum size to the detection threshold size, then we see that the time to grow to threshold size is simply $\frac{1}{k} ln \frac{1}{m}$.

Variability in the timing of recurrence in this scenario is created by variability in the amount of residuum left behind in the susceptible patients. We don't know exactly what distribution surgery or chemotherapy might leave behind in different patients, but we can set some likely constraints. The value of $m$ cannot be zero, since we are accounting for "cured" patients separately. It cannot be 1 because that would imply the tumor is detectable immediately after treatment. The average value of m is likely below 0.5, since otherwise one would expect more than half of susceptible patients to recur within a single doubling time.

A good choice for a distribution for $m$ to follow is the Beta Distribution, a continuous distribution defined on the interval [0,1], with a PDF of $x^{\alpha-1}(1-x)^{\beta-1}$. Choosing a Beta distribution with a first parameter of 1 and a second parameter > 2 produces a monotonically declining probability distribution function that has zero probability at x=1, but variable degrees of concavity between 0 and 1.

It is then straightforward to show that the cumulative distribution function for the distribution of $\frac{1}{k} ln \frac{1}{m}$, where $m$ follows such a Beta Distribution is simply $(1-e^{-kt})^{\beta}$, which means the survival function is $1-(1-e^{-kt})^{\beta}$. We may re-parametrize this expression in terms of the mean of the distribution, $\mu = \frac{1}{1+\beta}$, so that it becomes $1-(1-e^{-kt})^{-1+\frac{1}{\mu}}$, where $\mu$ may be understood as the average size of the tumor residua, relative to the detection threshold. Clearly $\mu$ <<1/2, or else average residua would reach threshold in a single doubling time. For modeling purposes, we shall assume $\mu$ < 1/3.

Finally, taking into account patients who are cured, we get a final survival function of

$$\rho + (1-\rho)\left(1-(1-e^{-kt})^{-1+\frac{1}{\mu}}\right), t > 0; \mu < 1/3$$

**Model 3: Variable tumor growth rate**

Here we consider the possibility that tumor residua remaining after treatment are all about the same size in all susceptible individuals, but the rate of re-growth varies across individuals, according to some distribution of the growth rate constant, $k$.

As in model 2, the time for a residuum of size $s$ to regrow at an exponential rate to a threshold size $\theta$ is $\frac{1}{k} ln \frac{\theta}{s}$ but now $k$ is the random variable rather than $s$. We thus replace $ln \frac{\theta}{s}$ with a single parameter, $\varphi$, that represents the log fold change in size between the threshold and starting sizes. Clearly $\theta/s > 1$, and most likely $\theta/s > 2$, otherwise tumors will re-grow to threshold size in less than one doubling time. Thus $\varphi > 0$ and most likely $\varphi > ln2$.

As for the distribution that $k$ should follow, we seek a flexible distribution that has support for only positive numbers, with a single peak and flexible degree of skewing toward large or small values. Data suggest that doubling times for a wide variety of human cancers are approximately log-normally distributed across patients with coefficients of variation typically less than 1 (Kay et al., 2019). Doubling time is simply $ln2/k$, so the distribution of times to reach threshold will be given by the distribution of $\varphi \times g/ln2$ where $g$ is log-normally distributed. The CDF for that is:

$$\frac{1}{2}\text{Erfc}\left[-\frac{ln\left[\sqrt{1+c^2}\,t\,\frac{ln2}{\kappa\varphi}\right]}{\sqrt{2\,ln[1+c^2]}}\right]$$

where $\kappa$ is the mean and $c$ is the coefficient of variation of the log normal distribution (note that $\kappa$ and $c$ refer to the actual mean and coefficient of variation of the distribution, and not those of the underlying normal distribution in terms of which the lognormal distribution is often parametrized).

We then define two lumped parameters, $\phi = \kappa\varphi/ln2$ and $\chi = \frac{\sqrt{ln[1+c^2]}}{\sqrt{2}}$. Note that $\phi$ is simply the time to reach threshold for a tumor residuum growing at the average rate, and $\chi$ is monotonically related to the coefficient of variation of the growth rate constant distribution. With these substitutions, the CDF becomes

$$1-\frac{1}{2}\text{Erfc}[-\frac{\chi^2+ln[t/\phi]}{2\chi}] = \frac{1}{2}\text{Erfc}[\frac{\chi^2+ln[t/\phi]}{2\chi}]$$

so that the full survival function becomes

$$\rho+(1-\rho)(\frac{1}{2}\text{Erfc}[\frac{\chi^2+ln[t/\phi]}{2\chi}]), t > 0$$

We note that the constraint that the coefficient of variation of the lognormal distribution be below 1 (Kay et al., 2019) constrains $\chi$ to be < 0.59.
